# Shotgun metagenomics and metabolomics reveal glyphosate alters the gut microbiome of Sprague-Dawley rats by inhibiting the shikimate pathway

**DOI:** 10.1101/870105

**Authors:** Robin Mesnage, Maxime Teixeira, Daniele Mandrioli, Laura Falcioni, Quinten Raymond Ducarmon, Romy Daniëlle Zwittink, Caroline Amiel, Jean-Michel Panoff, Fiorella Belpoggi, Michael N Antoniou

## Abstract

There is intense debate as to whether glyphosate can interfere with aromatic amino acid biosynthesis in microorganisms inhabiting the gastrointestinal tract, which could potentially lead to negative health outcomes. We have addressed this major gap in glyphosate toxicology by using a multi-omics strategy combining shotgun metagenomics and metabolomics. We tested whether glyphosate (0.5, 50, 175 mg/kg bw/day), or its representative EU commercial herbicide formulation MON 52276 at the same glyphosate equivalent doses, has an effect on the rat gut microbiome in a 90-day subchronic toxicity test. Clinical biochemistry measurements in blood and histopathological evaluations showed that MON 52276 but not glyphosate was associated with statistically significant increase in hepatic steatosis and necrosis. Similar lesions were also present in the liver of glyphosate-treated groups but not in the control group. Caecum metabolomics revealed that glyphosate inhibits the enzyme 5-enolpyruvylshikimate-3-phosphate (EPSP) synthase in the shikimate pathway as evidenced by an accumulation of shikimic acid and 3-dehydroshikimic acid. Levels of caecal microbiome dipeptides involved in the regulation of redox balance (γ-glutamylglutamine, cysteinylglycine, valylglycine) had their levels significantly increased. Shotgun metagenomics showed that glyphosate affected caecum microbial community structure and increased levels of *Eggerthella spp*. and *Homeothermacea spp..* MON 52276, but not glyphosate, increased the relative abundance of *Shinella zoogleoides*. Since *Shinella spp.* are known to degrade alkaloids, its increased abundance may explain the decrease in solanidine levels measured with MON 52776 but not glyphosate. Other glyphosate formulations may have different effects since Roundup® GT Plus inhibited bacterial growth *in vitro* at concentrations at which MON 52276 did not present any visible effect. Our study highlights the power of a multiomics approach to investigate effects of pesticides on the gut microbiome. This revealed the first biomarker of glyphosate effects on rat gut microbiome. Although more studies will be needed to ascertain if there are health implications arising from glyphosate inhibition of the shikimate pathway in the gut microbiome, our findings can be used in environmental epidemiological studies to understand if glyphosate can have biological effects in human populations.

**Graphical Abstract:** 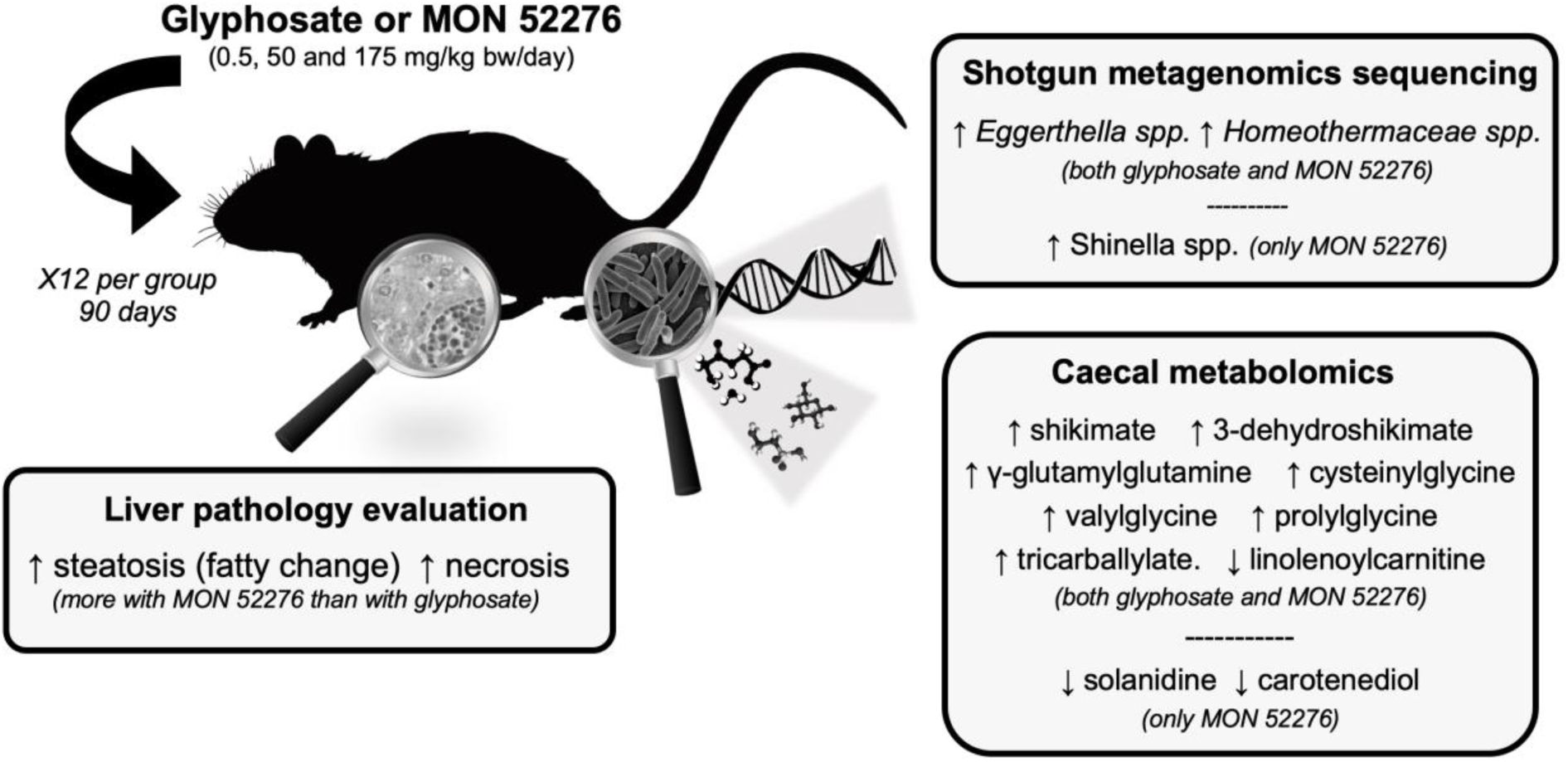

## Introduction

Glyphosate is the world’s most used herbicide ingredient (Benbrook 2016) and is one of the most frequently detected pesticide residues in foodstuffs (EFSA, 2017). This is due to the fact that it is frequently sprayed on crops (especially cereals such as oats and wheat) to accelerate ripening and facilitate harvest, or to clear weeds in the cultivation of glyphosate-tolerant genetically modified crops (Benbrook 2016; EFSA 2017).

Glyphosate acts as a weedkiller by inhibiting 5-enolpyruvylshikimate-3-phosphate synthase (EPSPS) in plants (Schonbrunn et al. 2001). The mode of action of glyphosate on EPSPS is to compete with phosphoenolpyruvate, which is condensed with shikimate-3-phosphate to form 5-enolpyruvylshikimate-3-phosphate (EPSP). The action of EPSPS is the penultimate step in the seven-step shikimate pathway leading to the biosynthesis of chorismate (Knaggs 2001). Although it is generally considered that the inhibition of aromatic amino acid synthesis is the main outcome of glyphosate effects on the shikimate pathway, chorismate is also a precursor for the biosynthesis of secondary metabolites including ubiquinone, menaquinone, lignans, tannins and flavonoids (Knaggs 2001).

As the shikimate pathway is absent in animal cells, including humans, glyphosate has been asserted to have a very high safety profile. However, the shikimate pathway also exists in some microorganisms. Since many microorganisms rely on EPSPS for aromatic amino acid biosynthesis, it has been found that glyphosate in combination with dicarboxylic acids (for example oxalate) acts as an antiparasitic agent and can potentially be used for the treatment of infections caused by protozoan parasites such as *Toxoplasma gondii, Plasmodium falciparum* and *Cryptosporidium parvum*. (U.S. Patent No 7771736 B2). By extrapolation, glyphosate was proposed to function as an antibiotic against shikimate pathway positive bacteria (U.S. Patent No 7771736 B2). In addition, glyphosate has been described to have fungicidal properties when sprayed in agricultural fields. For example, spraying Roundup tolerant genetically modified crops with a glyphosate-based herbicide (GBH) during their cultivation inhibited the growth of *Phakopsora pachyrhizi* and *Puccinia spp.* known to cause rust diseases (Feng et al. 2005). In another study of vineyard soil microorganisms, the presence of glyphosate residues was suggested to act as a selective fungicide favouring the growth of *Colletotrichum sp., Cunninghamella sp., Mortierella sp. and Scedosporium sp*. (Mandl et al. 2018). Despite the growing scientific evidence linking the presence of fungi in the human gut to health effects, no study has investigated if glyphosate can affect these fungal populations known as the gut ‘mycobiome’ (Huseyin et al. 2017).

Glyphosate is primarily excreted in faeces, which makes the content of the gastrointestinal tract the biological compartment exposed to the highest concentrations of this compound (Brewster et al. 1991). It is still unclear whether glyphosate is metabolised by the gut microbiome (Zhan et al. 2018). The main metabolite of glyphosate is aminomethylphosphonic acid (AMPA), which can be synthesised by some soil microorganisms through the glycine oxidase pathway, or by oxidation of the glyphosate carbon–phosphorus bond (Barrett and McBride 2005). Another biochemical pathway found in some soil microorganisms is the C-P lyase pathway, also called the sarcosine pathway, which cleaves the glyphosate C-P bond to phosphate and sarcosine (Hove-Jensen et al. 2014).

The gastrointestinal tract is colonised by a large collection of microorganisms including bacteria, archea, fungi, viruses, and small eukaryotes. Bacteria are physically separated from the gut epithelial lining by the inner mucus layer, which prevents epithelial colonisation and potentially enteric infection (Ducarmon et al. 2019). Gut microbiome chemical metabolism has a major influence on health (Mesnage et al., 2019). The large enzymatic repertoire carried by microorganisms in the gut confers on them the ability to influence the therapeutic effects of common drugs such as acetomiphen (Clayton et al., 2009) or digoxin (Haiser et al. 2013). A recent study of the relationship between the human faecal microbiome and the blood metabolome on 1004 human twin pairs showed that half of the blood metabolites (46%) were associated with microbial species and/or functional pathways (Visconti et al. 2019). As a consequence, changes in the human gut microbiome influences a wide range of clinical conditions such as Crohn’s disease, type 2 diabetes, obesity, and behavioural problems (Valdes et al., 2018).

It has been hypothesized that glyphosate may contribute to the development and progression of various human diseases by generating a selection pressure on some microbial communities in the human gut microbiome (Mesnage and Antoniou 2017). Although some studies have investigated the effects of glyphosate on the gut microbiome in rats (Lozano et al. 2018; Mao et al. 2018; Nielsen et al. 2018), cows (Riede et al. 2016), honey bees (Motta et al. 2018) or turtles (Kittle et al. 2018), there is still intense debate as to whether glyphosate interference with the shikimate pathway in microorganisms inhabiting the human gastrointestinal tract can be a source of negative health outcomes.

In order to address this knowledge gap in glyphosate toxicology, we have used a multi-omics strategy combining caecal microbiome shotgun metagenomics and metabolomics, to test whether glyphosate, or its representative EU commercial herbicide formulation MON 52276, has an effect on the rat gut microbiome. We took advantage of recent progress in high-throughput “omics technologies”, which are used to evaluate molecular composition and which predicts chemical mode of action (Taylor et al. 2018; Zampieri et al. 2018). Metabolomics is increasingly used to understand the function of the gut microbiome (Zierer et al. 2018). Combined with shotgun metagenomics sequencing techniques to identify and quantify the whole genomes from a larger range of micro-organisms (bacteria, fungi, viruses and protists), it has become possible to capture the metabolic activity of the gut microbiome. This strategy allowed us to demonstrate that glyphosate inhibits the shikimate pathway in the rat gut microbiome, causing the accumulation of shikimate pathway intermediates, which can be associated with changes in host metabolism and redox balance.

## Material and methods

### Experimental animals

The experiment was conducted on young adult female Sprague-Dawley rats (8 weeks of age at the start of treatment) in accordance with Italian law regulating the use and humane treatment of animals for scientific purposes (Decreto legislativo N. 26, 2014. Attuazione della direttiva n. 2010/63/UE in materia di protezione degli animali utilizzati a fini scientifici. – G.U. Serie Generale, n. 61 del 14 Marzo 2014). Before commencing the experiment, the protocol was examined by the animal-welfare body for approval. The protocol of the experiment was authorised by the ad hoc commission of the Italian Ministry of Health (authorization N. 447/2018-PR).

Female Sprague-Dawley rats were generated in-house at the Cesare Maltoni Cancer Research Center, Ramazzini Institute, (Bentivoglio, Italy) following an outbreeding programme, and were subjected to ear punch marking using the Jackson Laboratory system for the purposes of identification. Female animals were chosen in order to make our results comparable to our previous studies where we showed that the long-term exposure to Roundup® GT plus was associated with the development of fatty liver disease (Mesnage et al. 2015; Mesnage et al. 2017). Animals were randomised after weaning in order to have at most one sister per litter in each group. Homogeneous body weight within the different groups was also ensured. Animals of 6 weeks of age were acclimatised for two weeks before the start of the experiment. Rats were housed in polycarbonate cages (41×25×18 cm) with stainless steel wire tops and a shallow layer of white wood shavings as bedding. The animals were housed in the same room, 3 per cage, maintained at a temperature of 22±3°C and relative humidity of 50±20%. Artificial lighting was provided on a 12-hour light/dark cycle. The cages were periodically rotated on their racks to minimise effects due to cage position on animals.

### Treatments

Seven groups of 12 female Sprague-Dawley rats of 8 weeks of age were treated for 90 days. They received *ad libitum* a rodent diet supplied by SAFE (Augy, France). The feed was analysed to identify possible contaminants or impurities by Eurofins (Germany) and the Institut Scientifique d’Hygiène et d’Analyse (ISHA, Champlan, France) (Supplementary Material 1). Nutrients, pesticides, dioxins, mycotoxins, isoflavones, heavy metals, and polychlorinated biphenyls, as well as some microorganisms, were measured. Glyphosate was not detected (limit of quantification 0.01 mg/kg). The only pesticide residue detected was piperonyl butoxide found at a concentration of 0.034 (± 0.017) mg/kg. Glyphosate and MON 52276 (at the same glyphosate equivalent dose) were administered via drinking water to give a daily intake of 0.5 mg, 50 mg and 175 mg/kg body weight per day (mg/kg bw/day), which respectively represent the EU acceptable daily intake (ADI), the EU no-observed adverse effect level (NOAEL) and the US NOAEL (European Food Safety 2015). Glyphosate was purchased from Merck KGaA (Sigma Aldrich®, Gillingham, Dorset, UK) with purity ≥ 95%. MON 52276 was sourced from the Italian market as Roundup Bioflow (commercial name). MON 52276 is the formulated GBH used for the EU’s review for market registration, and sold under different trade names in Europe (Roundup Pro Biactive in Ireland, Roundup Extra 360 or Roundup Star 360 in France, Roundup BioFlow in Italy, Roundup Ultra in Belgium and Austria, or Roundup Bio in Finland) (Mesnage et al. 2019). Tap water from the local water supplier was administered in glass bottles *ad libitum*. Drinking water was discarded and the bottles were cleaned and refilled daily. The doses of MON 52776 and glyphosate for experimental groups were calculated every week on the basis of mean body weights and mean water consumption in order to achieve the desired dosage.

### Clinical observations

Animals were checked for general status three times a day, seven days a week, except for non-working days when they were checked twice. Status, behaviour and clinical parameters of experimental animals were checked weekly starting from 2 weeks before the start of the treatments until the end of the experiment (13 weeks of treatment). The daily water and food consumption per cage were measured before the start of the experiment, and weekly for the entire 13-week duration of the treatment. Before final sacrifice and after approximately 16 hours in a metabolic cage, total water consumption was registered for each animal. Body weight of experimental animals was measured before the start of the treatment, and then weekly for 13 weeks.

### Evaluation of pathologies

All sacrificed animals were subjected to complete necropsy. Liver and kidneys were alcohol-fixed, trimmed, processed and embedded in paraffin wax. Sections of 3-6 μm were cut for each specimen of liver and kidneys and stained with haematoxylin and eosin. All slides were evaluated by a pathologist and all lesions of interest were further assessed by a second more senior pathologist.

The histopathological nomenclature of lesions adopted was harmonised with international criteria. In particular, the non-neoplastic lesions were classified according to the international nomenclature INHAND (International Harmonization of Nomenclature and Diagnostic Criteria) and RITA (Registry of Industrial Toxicology Animal Data). Incidence of non-neoplastic lesions was evaluated with a Fisher’s exact test (one and two-tailed; one-sided results were also considered, since it is well established that only an increase in the incidences can be expected from the exposure, and incidences in the control group are almost always 0). The analysis of linear trend, for incidence of pathological lesion, was obtained using the Cochran-Armitage trend test (OECD 2011; Shockley and Kissling 2018).

### Biochemistry

Prior to sacrifice, animals were anesthetised by inhalation of CO_2_/O_2_ (70% and 30% respectively), and approximately 7.5 ml of blood was collected from the *vena cava*. Serum biochemistry was performed at IDEXX BioAnalytics (Stuttgart, Germany), a laboratory accredited to ISO 17025 standards. Sodium and potassium levels were measured by indirect potentiometry. Albumin was measured by a photometric Bromocresol green test. Alkaline phosphatase ALP was measured by IFCC with the AMP-buffer method, glucose by Enzymatic UV-Test (Hexokinase method), cholesterol by Enzymatic colour test (CHOD-PAP), blood urea nitrogen by enzymatic UV-Test, gamma-glutamyl-transferase by Kinetic colour test International Federation of Clinical Chemistry (IFCC), aspartate and alanine aminotransferase by kinetic UV-test (IFCC+ pyridoxal-5-phosphate), creatinine by kinetic colour test (Jaffe’s method), lactate dehydrogenase (LDH) by the IFCC method, and triglycerides using an enzymatic colour test (GPO-PAP) on a Beckman Coulter AU 480 instrument.

### Metabolomics analysis

Metabolon Inc. (Durham, NC, USA) was contracted to conduct the metabolomics analysis. Faecal samples were prepared using the automated MicroLab STAR® system from Hamilton Company. Proteins were precipitated with methanol under vigorous shaking for 2 min (Glen Mills GenoGrinder 2000), followed by centrifugation. Samples were placed briefly on a TurboVap® (Zymark) to remove the organic solvent. The sample extracts were stored overnight under nitrogen before preparation for analysis. The resulting extract was analysed on four independent instrument platforms: two different separate reverse phase ultra-high performance liquid chromatography-tandem mass spectroscopy analysis (RP/UPLC-MS/MS) with positive ion mode electrospray ionisation (ESI), a RP/UPLC-MS/MS with negative ion mode ESI, as well as a by hydrophilic-interaction chromatography (HILIC)/UPLC-MS/MS with negative ion mode ESI.

All UPLC-MS/MS methods utilised a Waters ACQUITY ultra-performance liquid chromatography (UPLC) and a Thermo Scientific Q-Exactive high resolution/accurate mass spectrometer interfaced with a heated electrospray ionization (HESI-II) source and Orbitrap mass analyser operated at 35,000 mass resolution. The sample extract was dried and then reconstituted in solvents compatible to each of the four methods used (Evans et al., 2014). Each reconstitution solvent contained a series of standards at fixed concentrations to ensure injection and chromatographic consistency. One aliquot was analysed using acidic positive ion conditions, chromatographically optimised for more hydrophilic compounds. In this method, the extract was gradient eluted from a C18 column (Waters UPLC BEH C18-2.1×100 mm, 1.7 µm) using water and methanol, containing 0.05% perfluoropentanoic acid (PFPA) and 0.1% formic acid (FA). Another aliquot was also analysed using acidic positive ion conditions, chromatographically optimised for more hydrophobic compounds. In this method, the extract was gradient eluted from the same aforementioned C18 column using methanol, acetonitrile, water, 0.05% PFPA and 0.01% FA and was operated at an overall higher organic content. Another aliquot was analysed using basic negative ion optimized conditions using a separate dedicated C18 column. The basic extracts were gradient eluted from the column using methanol and water, with 6.5mM ammonium bicarbonate at pH 8. The fourth aliquot was analysed via negative ionisation following elution from a HILIC column (Waters UPLC BEH Amide 2.1×150 mm, 1.7 µm) using a gradient consisting of water and acetonitrile with 10mM ammonium formate, pH 10.8. The MS analysis alternated between MS and data-dependent MSn scans using dynamic exclusion. The scan range varied slightly between methods but covered 70-1000 m/z. Raw data was extracted, peak-identified and QC processed using Metabolon’s hardware and software as previously described (Mesnage et al. 2018).

### Shotgun metagenomics

At the time of sacrifice, two vials of 100 mg caecal content were collected and stored to perform gut microbiome evaluation. DNA was extracted from 100 mg caecum content using the Quick-DNA Fecal/Soil Microbe Miniprep Kit (ZymoResearch) with minor adaptations from manufacturer instructions. Adaptations were: 1. bead beating was performed at 5.5 m/s for three times 60 seconds (Precellys 24 homogeniser, Bertin Instruments) and 2. 50 µl elution buffer was used to elute the DNA following which the eluate was run over the column once more to increase DNA yield. One negative control (no sample added) and one positive control (ZymoBIOMICS Microbial Community Standard, ZymoResearch) were taken along during the DNA extraction procedures and subsequently sequenced. DNA was quantified using the Qubit HS dsDNA Assay kit on a Qubit 4 fluorometer (Thermo Fisher Scientific).

Shotgun metagenomics was performed under contract by GenomeScan (Leiden, The Netherlands). The NEBNext® Ultra II FS DNA module (cat# NEB #E7810S/L) and the NEBNext® Ultra II Ligation module (cat# NEB #E7595S/L) were used to process the samples. Fragmentation, A-tailing and ligation of sequencing adapters of the resulting product was performed according to the procedure described in the NEBNext Ultra II FS DNA module and NEBNext Ultra II Ligation module Instruction Manual. The quality and yield after sample preparation was measured with the Fragment Analyzer. The size of the resulting product was consistent with the expected size of approximately 500-700 bp. Clustering and DNA sequencing using the NovaSeq6000 was performed according to manufacturer’s protocols. A concentration of 1.1 nM of DNA was used. NovaSeq control software NCS v1.6 was used.

Shotgun metagenomics datasets were analysed with Rosalind, the BRC/King’s College London high-performance computing cluster. First, data was pre-processed using preprocessing package v0.2.2 (https://anaconda.org/fasnicar/preprocessing). In brief, this package concatenates all forward reads into one file and all reverse reads into another file, then uses trim_galore to remove Illumina adapters, trim low-quality positions and unknown position (UN) and discard low-quality (quality <20 or >2 Ns) or too short reads (< 75bp) removes contaminants (phiX and rat genome sequences), and ultimately sorts and splits the reads into R1, R2, and UN sets of reads. After pre-processing, a total of 20.7 ± 4.9 million paired-end non-rat cleaned reads remained.

Remaining reads were then aligned to a bacterial gene catalogue from the Sprague-Dawley rat gut microbiome using Bowtie2 version 2.2.5 (Pan et al. 2018). A total of 23.5 ± 6.8% of the reads aligned to this catalogue. Count data was then extracted from the BAM files using SAMtools (v1.4). We also complemented the taxonomic composition analysis by using the gene marker-bbased tool IGGsearch (Nayfach et al. 2019) because the classification of contigs at the species level in the Sprague-Dawley rat gut microbiome gene catalogue was quite poor. A total of 90.6% of the contigs detected in our metagenome datasets were not annotated at the species level in the rat gut microbiome catalogue. This analysis was done using the iggdb_v1.0.0_gut database and default parameters (Nayfach et al. 2019). ShortBRED was also used to quantify protein families from the shikimate pathway with default parameters (Kaminski et al. 2015).

### *In vitro* study of bacterial growth

A stock solution of glyphosate was made in water and used to inoculate bacterial growth medium. Roundup® GT plus (GT^+^) and MON 52276 at 450 g/L and 360 g/L respectively, were directly diluted in bacterial broth. *Lactobacillus rhamnosus* strains were provided by the UCMA’s (Université de Caen Microbiologie Alimentaire) culture collection ID 2927, 2929, 2933 and 5164. These strains were used because *Lactobacillus rhamnosus* is prevalent in the human gut with relevant health-promoting properties as a probiotic (Capurso 2019). The broth dilution method was used to determine how pesticides can modify bacterial growth under aerobic conditions. Bacteria from overnight cultures were re-suspended to an OD_600nm_ = 0.3 and further diluted 1,000-fold in De Man, Rogosa and Sharpe (MRS) agar without peptone for *L. rhamnosus* to obtain approximately 10^5^ CFU/mL as confirmed in each experiment by plating the cell suspension on agar plates and incubated aerobically at 37°C. Positive and negative controls were included in each experiment. Plates were inspected after 24h and 48h. All experiments were performed in triplicate.

### Statistical analysis

Statistical analysis was performed in R version 3.6.1. The metabolome data analysis was performed using MetaboAnalyst 4.0 (Chong et al. 2018). Peak area values were median scaled, log transformed, and any missing values imputed with sample set minimums, both on a per biochemical basis as previously described (Mesnage et al. 2018). Log-transformed data was analysed using an ANOVA test adjusted for multiple comparisons with Fisher’s least significant difference in order to determine if there was a difference in mean value between the different treatment groups. For the shotgun metagenomics, a non-parametric approach on the taxonomic composition data was used since gut metagenomics datasets are typically zero-inflated (Knight et al. 2012). A total sum scaling normalisation was performed to account for differences in sequencing depth. Kruskal-Wallis test adjusted for multiple comparisons using the Benjamini–Hochberg procedure was performed with R in-house functions to assess differentially abundant taxonomic groups. Data analysis using count data extracted from the Sprague-Dawley gut microbiome catalogue was performed using DESeq2 (version 1.26.0) (Love et al. 2014) to detect the differentially abundant ontology KO (KEGG Orthology) terms. Pathway analysis was performed by mapping the different KO term to their KEGG pathway using the database available online (https://www.genome.jp/kegg/, downloaded on 24^th^ of October, 2019). Pathway enrichment analysis was performed using clusterProfiler (Yu et al. 2012). Statistical analyses of *in vitro* tests on bacterial growth were performed using GraphPad Prism version 8.0.1 (GraphPad Software, Inc, CA). Differences between treatment groups at different concentrations are performed using Kruskal-Wallis with Dunn’s multiple comparison post-test.

## Results

The primary aim of this study was to determine the effects of glyphosate on the rat gut microbiome. However, as commercial GBH products are known to be more toxic than glyphosate alone because they contain other potentially toxic compounds (Mesnage et al. 2019), we deemed it important to compare the effects of glyphosate to a representative EU Roundup formulation (MON 52276) administered at the same glyphosate equivalent dose. The doses tested were 0.5, 50 and 175 mg/kg bw/day glyphosate. These were chosen since they respectively represent the EU ADI, the EU NOAEL and the US NOAEL and are thus of regulatory relevance.

### Blood clinical biochemistry measurements and histopathological evaluation

No significant differences were observed in water consumption (Figure 1A), feed consumption (Figure 1B) and mean body weight (Figure 1C), except for the group administered with the highest dose of MON 52276 (175 mg/kg bw/day glyphosate equivalent dose), which had a reduced water consumption. All treated and control animals were subjected to a histopathological evaluation of the kidney and liver. No lesions were detected in liver of control animals whereas a low frequency (2/8 animals) of pelvic mineralisation, inflammation and epithelial pelvic necrosis was observed in their kidneys (Figure 1D, white shaded columns). Contrastingly, a dose-dependent and statistically significant increase in the incidence of liver lesions (fatty liver changes, necrosis) was observed in rats treated with MON 52276 (Figure 1D, orange shaded columns). An increase in liver lesions was also detected in animals treated with glyphosate, although this did not reach statistical significance (Figure 1D, blue shaded columns). No lesions were detected in the control group (Figure 1D, white shaded columns). Blood clinical biochemistry only revealed increased creatinine levels in the groups treated with glyphosate but not with MON 52276 (Figure 1E). Considering the limited time of exposure (90 days), and the limited number of animals, it is possible that this non-statistically significant alterations in liver and kidney structure could be due to the glyphosate or MON 52276 exposure.

**Figure 1.**
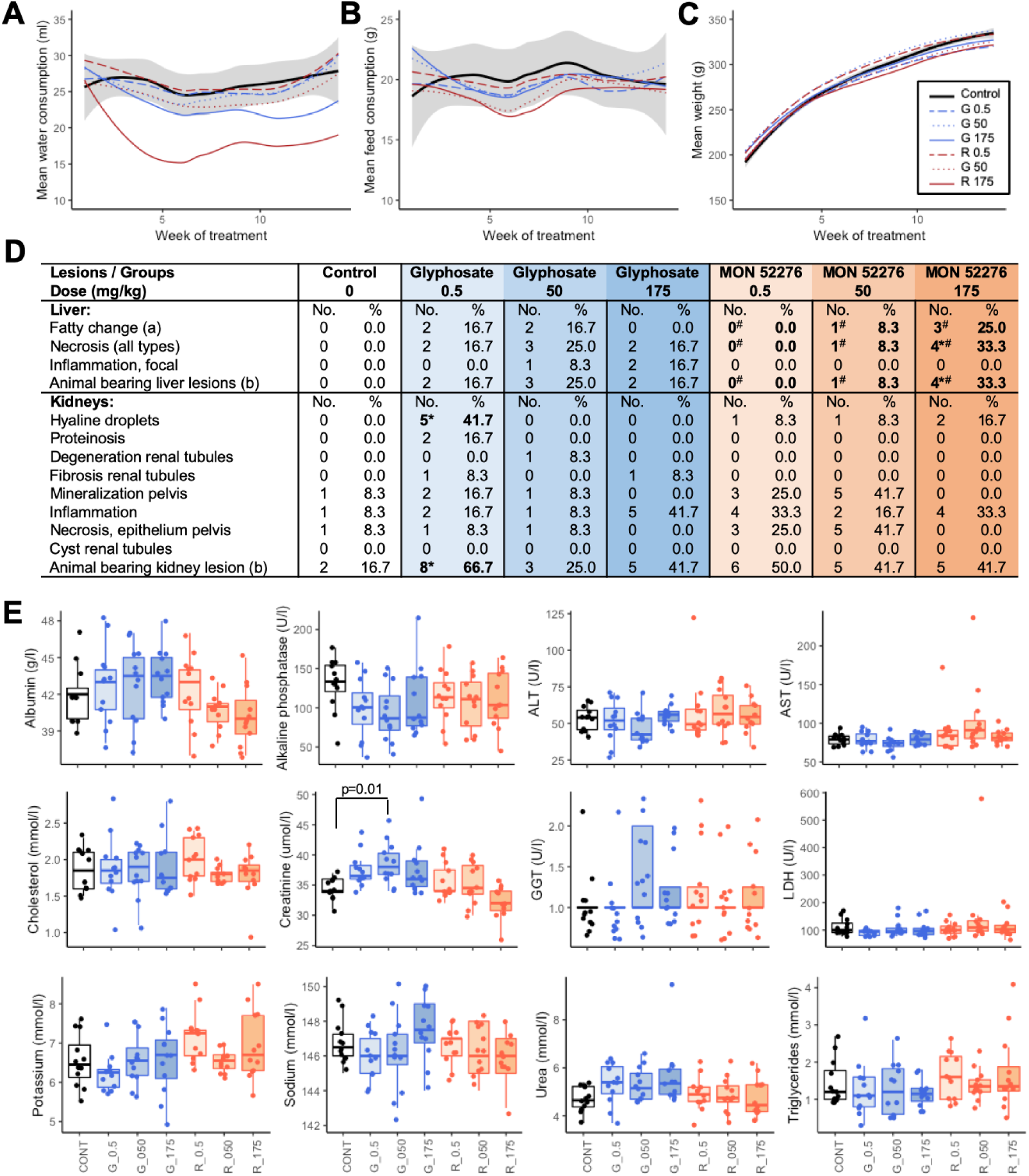
Subchronic toxicity of glyphosate and Roundup MON 52276 administered in drinking water to adult female Sprague-Dawley rats for 90 days. No differences in either water **(A)** and food **(B)** consumption, or in body weights **(C)** were detected, expect for water consumption of animals treated with Roundup at the highest dose, which was lower compared to controls. Histopathological evaluation of liver and kidneys of all treated and control animals **(D)** revealed increased incidences of lesions in the livers of animals treated with Roundup: (a) fatty change includes from mild to severe lesions or associated to necrosis; (b) one animal could bear more than one lesion. *Statistically significant (P≤0.05) using Fisher Exact test (one-tailed test); ^#^ p-values (P≤0.01) associated with the Cochran-Armitage test for trend. **(E)** Serum clinical biochemistry evaluation at the end of the treatment period only revealed minor changes (One-way ANOVA with post-hoc Tukey HSD).

### Caecum metabolomics

A total of 744 metabolites were measured in the caecal content of rats exposed to glyphosate and MON 52276. A large number of metabolites were affected in a dose-dependent manner (Figure 2B). The levels of twelve metabolites were statistically significantly altered when assessed using an ANOVA test adjusted for multiple comparisons with Fisher’s least significant difference (adjusted p value < 0.05). The most striking effect was an accumulation of shikimate and 3-dehydroshikimate, which are metabolites upstream of the reaction catalysed by EPSPS in the shikimate pathway (Figure 2A). These metabolites were undetectable in samples from untreated rats (Figure 2C). This indicates that glyphosate inhibits the EPSPS enzyme in the rat gut microbiome.

**Figure 2.**
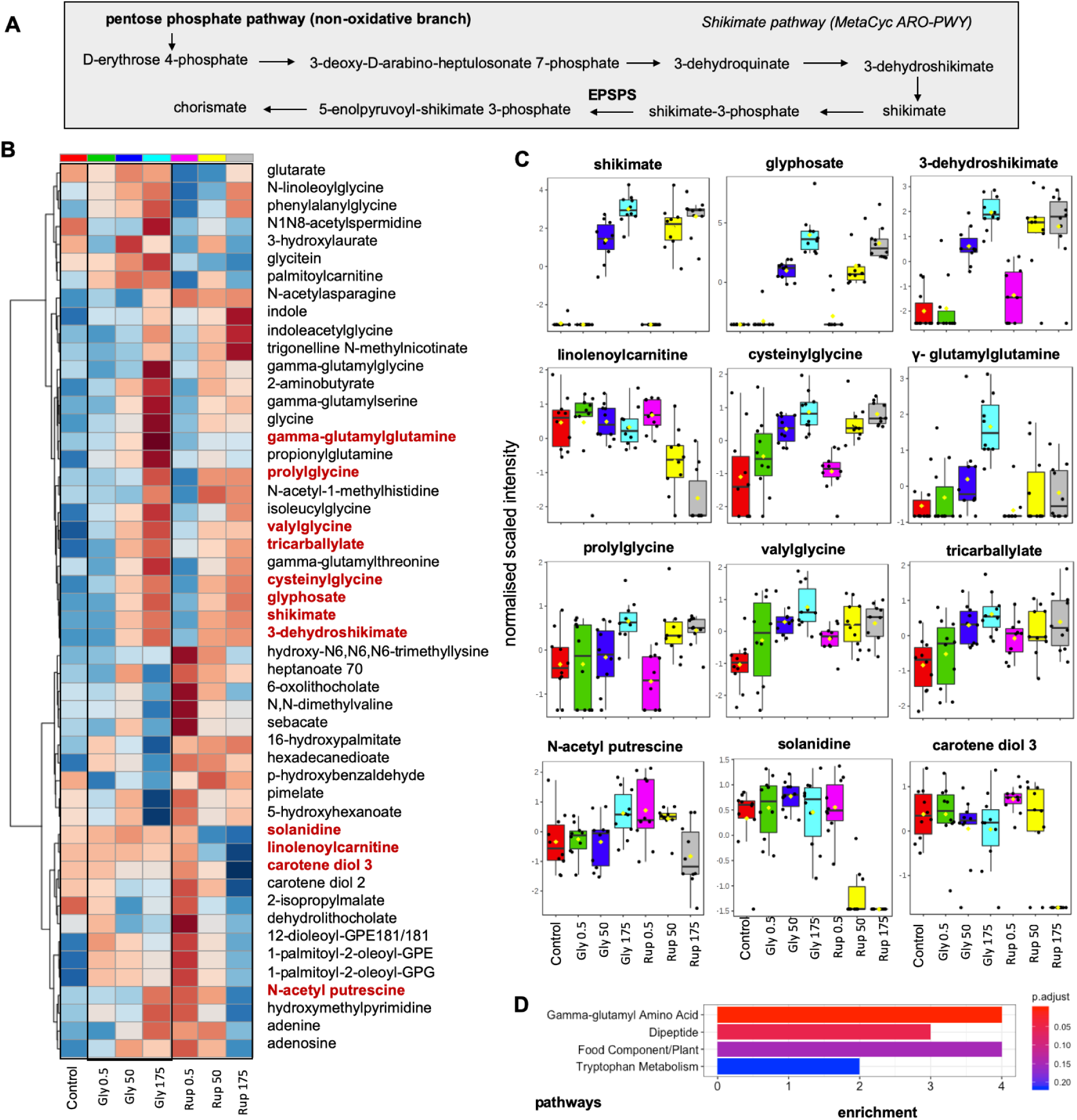
Glyphosate inhibits EPSPS in the rat gut microbiome. Caecal metabolomics reveals a dose-dependent accumulation of shikimate pathway intermediates reflective of EPSPS inhibition in rats exposed to increasing doses of glyphosate and its commercial formulation MON 52776. **A.** The shikimate pathway is the main target of glyphosate by inhibiting the enzyme 5-enolpyruvylshikimate-3-phosphate synthase (EPSPS).**B.** Hierarchical clustering of the 50 metabolites with the lowest p-values (adjusted p-value < 0.05 in red). **C.** Dot plots of statistically significant changes in the abundance for 12 metabolites show a dose dependent response to MON 52276 or/and glyphosate. **D.** Pathway enrichment analysis shows over-representation of gamma-glutamyl amino acid metabolism. The x-axis indicates enrichment as a fold-change.

Dipeptides involved in the regulation of redox balance (γ-glutamylglutamine, cysteinylglycine, valylglycine) were also found to have their levels significantly increased by MON 52276 and glyphosate treatments in a dose-dependent manner. Fold changes for these compounds generally ranged between 2 and 3, reaching a maximum of 4.4 for γ-glutamylglutamine at the highest dose of glyphosate. Pathway enrichment analysis also confirmed that glyphosate affected the level of metabolites involved in gamma-glutamyl amino acid metabolism (Figure 2D). Although most changes were very similar between the groups exposed to glyphosate and the groups exposed to MON 52276, additional changes were detected in the latter suggesting that formulated products induce additional effects over and above that of glyphosate alone. The most striking example was a decrease of solanidine and carotenediol levels to an extent that they became undetectable at the highest dose of MON 52276.

### Shotgun metagenomics analysis

We then performed a shotgun metagenomics analysis of the caecum microbiome. Taxonomic analysis at the phylum level found no statistically significant differences in relative abundance (Supplementary Material 2). A total of 1270 species-level operational taxonomic units were identified. No species were detected in a negative extraction control, which was included to ensure that no bacterial contamination was introduced by laboratory reagents and procedures. All the species present in both the ZymoBIOMICS Microbial Community Standard control and the IGGsearch database were detected in this taxonomy analysis. A total of 75% of the total abundance was assigned to 25 species dominated by *Fournierella sp.* (9.6%), *Lactobacillus murinus* (7.2%) and *Christensenellales sp.* (6.8%) (Figure 3A). Kruskal Wallis non-parametric analysis revealed that six species had their abundance altered by treatment with glyphosate or MON 52276 (Benjamini-Hochberg adjusted p-value < 0.05). Some species such as *Eggerthella* isolate HGM04355 and *Homeothermaceae* isolate HGM05183 were increased by treatment with the highest doses of both glyphosate and MON 52276. (Figure 3B). Interestingly, *Shinella zoogleoides* was increased by the treatment with MON 52276 from the lowest dose tested whilst no effect was observed with glyphosate.

**Figure 3.**
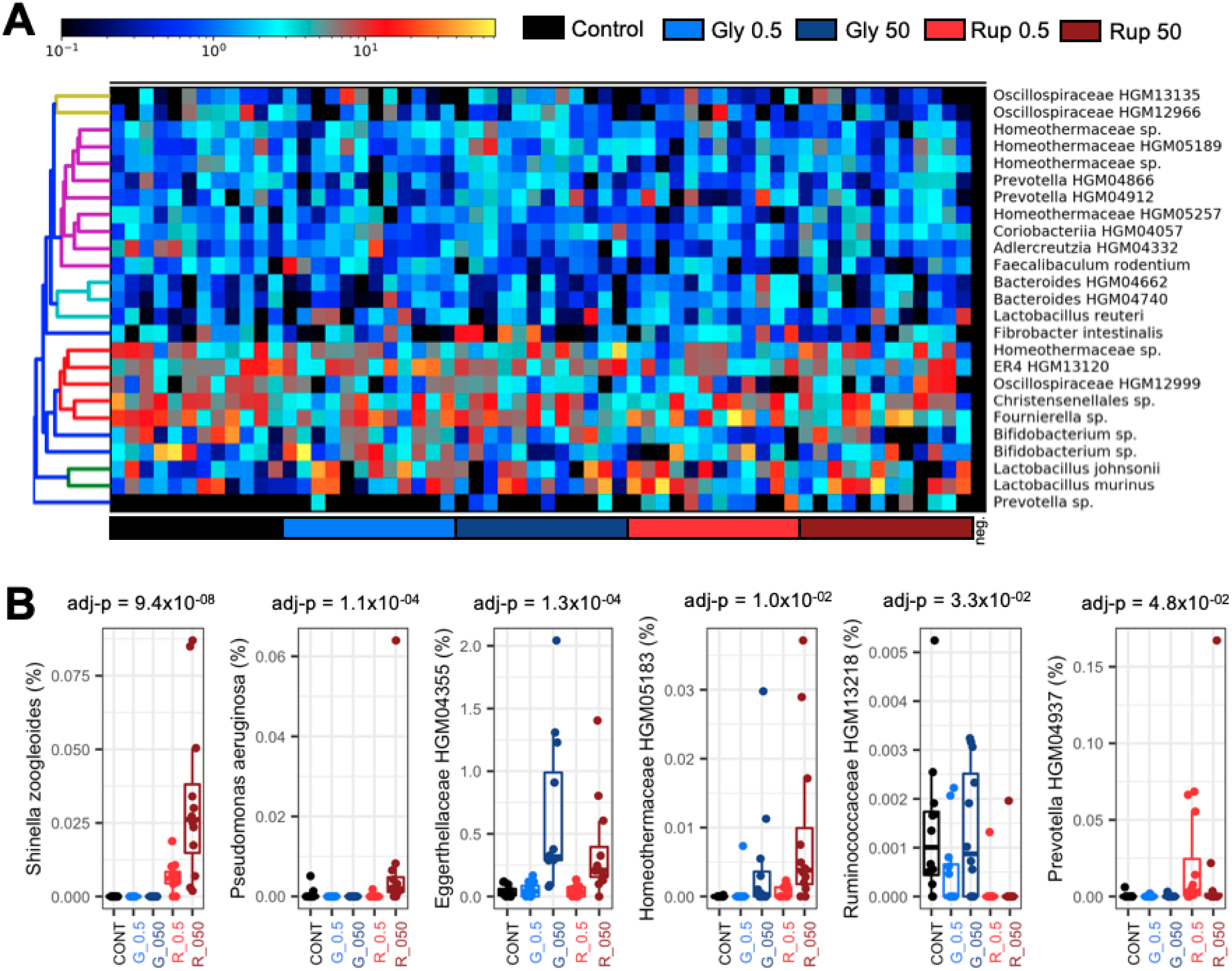
Shotgun metagenomics of glyphosate effects on the rat caecal microbiome. **A.** Top 25 species in the rat caecal microbiome using the marker gene-based tool IGGsearch **B.** Scatter plots displaying individual changes in relative abundance for the 6 species found to be affected by treatment with MON52276 and glyphosate (Kruskal–Wallis with Benjamini-Hochberg adjusted p-values).

We also performed a pathway analysis using the KO (KEGG Orthology) assignments provided in the Sprague-Dawley rat gut microbiome gene catalogue. Differential abundance estimates using DeSeq2 revealed that 1881 genes (out of 1,981,145 identified) had their abundance statistically significantly altered by exposure to glyphosate and MON 52776 as compared to the control group. However, differences in abundance between individual animals within a group were generally greater than the effect of test compounds, which limits the conclusions that can be drawn on the biological relevance of the observed statistically significant differences. Nevertheless, in order to understand the potential biological significance of these differences, a KEGG pathway enrichment analysis was performed. A large number of pathways were affected (Figure 4A). The most affected genes were related to the two-component regulatory system, which serves as a stimulus-response mechanism to allow bacteria to adapt to changing environments. It should be noted that the gene with the lowest p-value is annotated as a gamma-glutamyltranspeptidase (Figure 4B, first panel). We ultimately used ShortBRED to quantify the abundance of the EPSPS enzyme in the caecal metagenomes using a set of 10,809 markers created from the protein sequences of the InterPro family IPR006264. The most significant difference after a Kruskal–Wallis statistical test was an increase in the gene count for the 3-phosphoshikimate 1-carboxyvinyltransferase protein R7B7W1 from *Eggerthella sp. CAG:298*, which was detected at the higher dose (50mg/kg bw/day) of both glyphosate and MON 52276 (Figure 4C).

**Figure 4.**
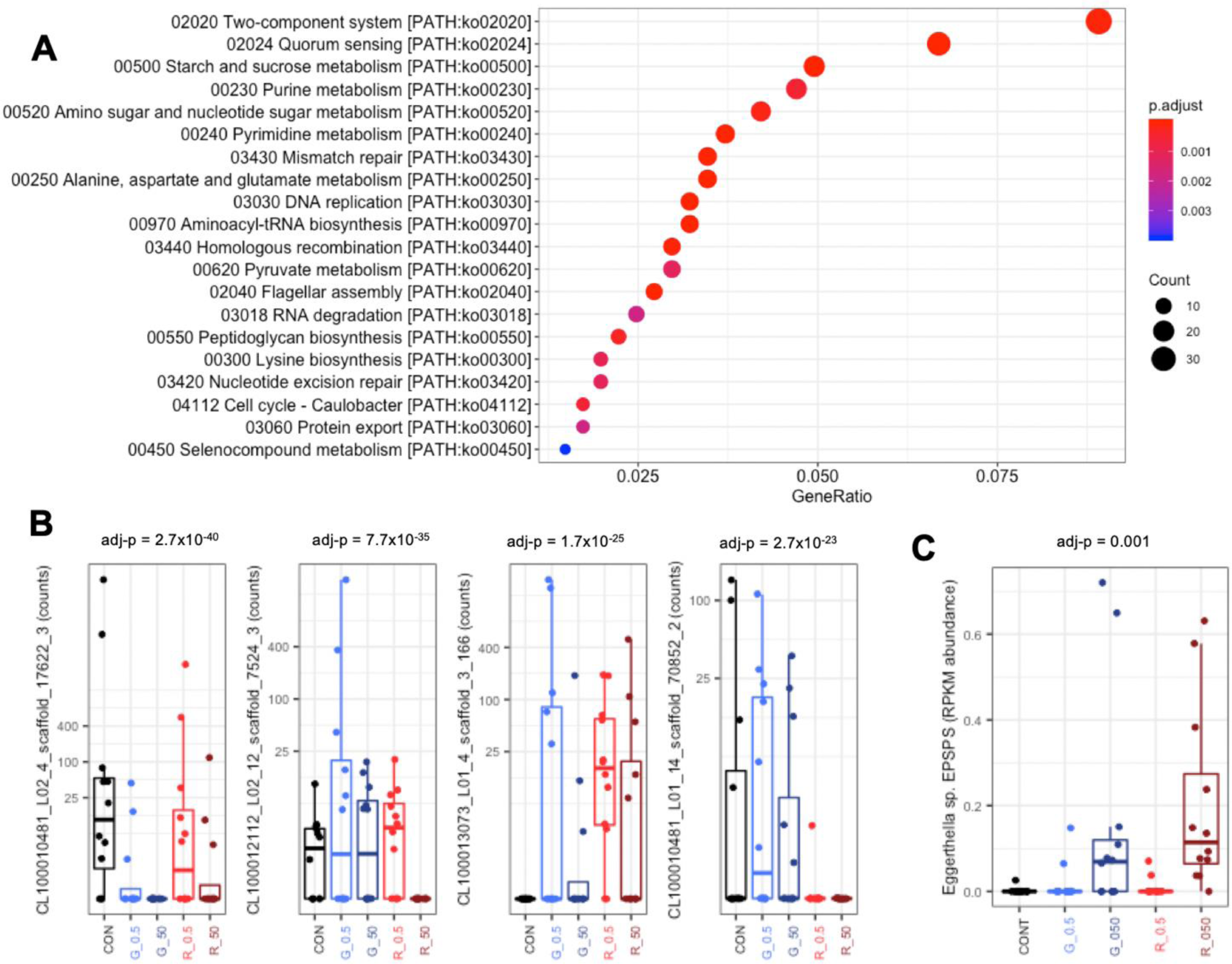
Over-representation (enrichment) of KEGG pathways determined using the gene catalogue of the Sprague-Dawley gut microbiome. **A**. Abundance of genes was considered to be statistically significant when their adjusted p-values were below 0.05 using DESeq2. **B**. Visual inspection of dispersion in abundance of the most significant contigs shows that differences in abundance between individual animals within a group were almost always greater than the effect of test compounds. **C.** ShortBRED analysis of InterPro family IPR006264 reveals that the 50mg/kg bw/day dose of glyphosate and MON52276 caused an increased abundance of the gene counts for the 3-phosphoshikimate 1-carboxyvinyltransferase protein R7B7W1 from *Eggerthella sp. CAG:298.*

### Growth inhibition in vitro

Since glyphosate formulated products are known to have different effects depending on their co-formulant composition, we compared the growth inhibition properties of glyphosate and MON 52276 to Roundup® GT Plus on four different strains of *Lactobacillus rhamnosus.* We observed that the growth of every bacterial strain was similarly inhibited by MON 52276 and glyphosate (Figure 5), which is consistent with the results obtained in the rat sub-chronic toxicity experiments. Interestingly, Roundup® GT Plus inhibited bacterial growth at concentrations at which glyphosate and MON 52276 did not present any visible effects. Another interesting finding is that glyphosate stimulated the growth of the strain LB2 whilst other strains were not affected (Figure 5).

**Figure 5.**
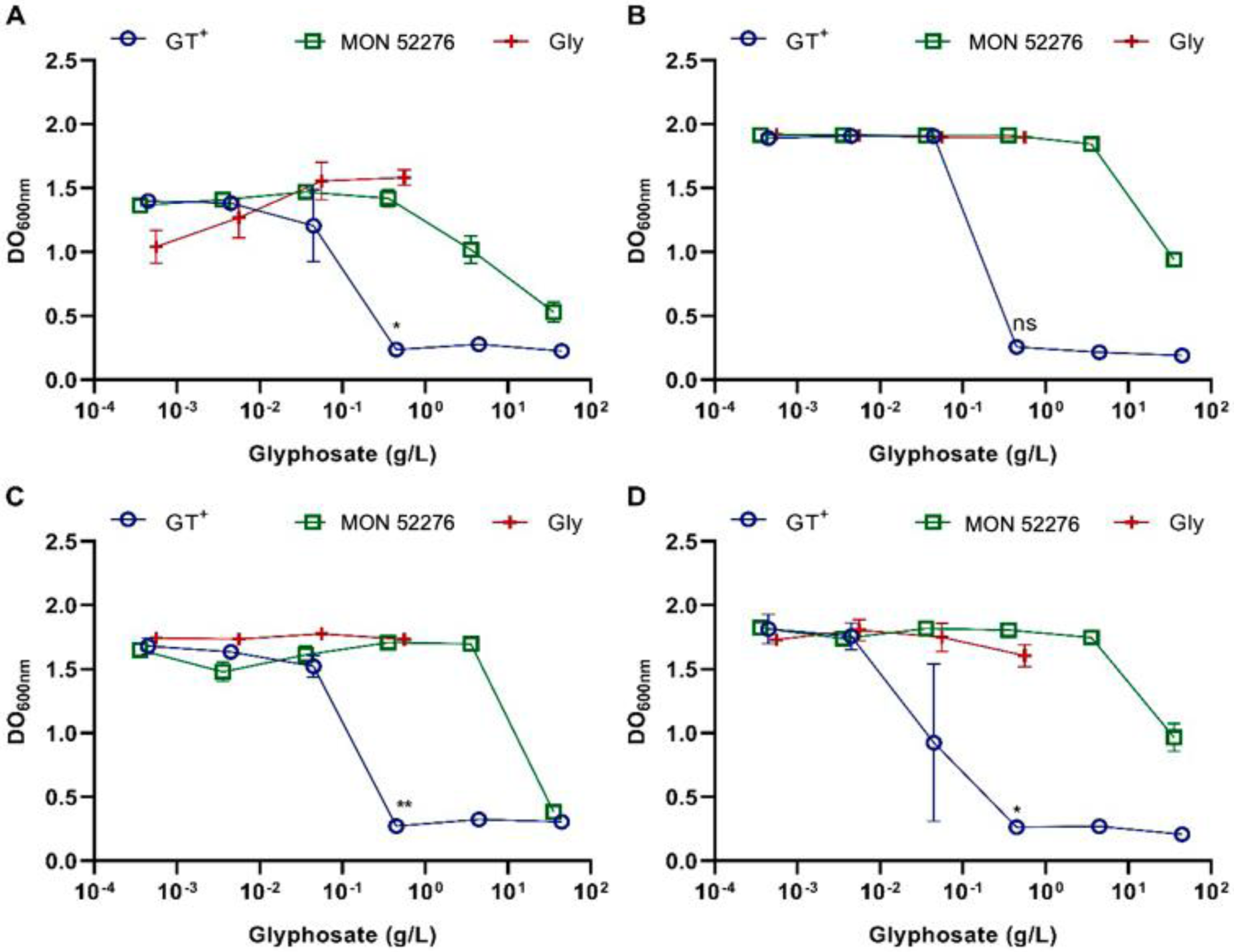
In vitro bacterial growth of four different *Lactobacillus rhamnosus* strains exposed for 48h to glyphosate alone or to different commercial formulations (Roundup GT^+^ or MON 52276). Bacterial growth of the strain *Lactobacillus rhamnosus* LB2 **(A)**, LB3 **(B)**, LB5 **(C)** and LB7 **(D)** was more inhibited when exposed to GT^+^ than to MON 52276.

Collectively, shotgun metagenomics revealed that glyphosate and MON52276 altered rat gut microbiome composition. This was visible from the lowest dose tested (0.5mg/kg bw/day), corresponding to the EU regulatory permitted ADI.

## Discussion

The effects of glyphosate on the gut microbiome are widely debated with divergent outcomes in studies performed to date. Some studies have indicated that glyphosate can cause alterations in the population of gut microbiota while others have not (Tsiaoussis et al. 2019). One major reason for the discrepancies regarding glyphosate effects on the gut microbiome, may be due to the lack of studies using advanced molecular profiling analytical techniques. In order to address this possibility, we have used a combination of shotgun-metagenomics and metabolomics to investigate the caecum microbiota of rats exposed to increasing doses of glyphosate and MON 52276. As a result, we were able to identify the first biomarker of glyphosate effects on the rat gut microbiome. We demonstrate for the first time that glyphosate inhibits the shikimate pathway in the rat caecum microbiome and is associated with changes in bacterial community structure. Although the rat gut microbiome is substantially different from that of humans, we anticipate that our findings will be of relevance for human physiology since the bacterial species inhabiting the human gastrointestinal tract have been found to be sensitive to glyphosate-mediated EPSPS inhibition (Tsiaoussis et al. 2019). However, epidemiological studies will be necessary to ascertain if the doses of glyphosate to which human populations are typically exposed are sufficient to change gut microbiome metabolism.

We found that glyphosate inhibition of EPSPS enzymes in the caecum microbiome led to an accumulation of intermediates of the shikimate pathway (Figure 2). This mechanism also leads to increases in shikimic acid in soil microorganisms (Aristilde et al. 2017). Shikimic acid can have multiple biological effects. Shikimate-rich plants such as *Illicium verum* Hook. f. (Chinese star anise) have been traditionally used to treat skin inflammation and stomach aches (Rabelo et al. 2015). Shikimic acid is a plant polyphenolic compound known to protect against oxidative stress (Rabelo et al. 2015), and has anti-platelet and anti-thrombogenic effects (Veach et al. 2016). Other studies have shown that shikimate can cause a dose-dependent activation of the aryl hydrocarbon receptor (AhR), a ligand-activated transcription factor having important roles in multiple tissues including in the mucosal immune system (Sridharan et al. 2014). However, deleterious or beneficial effects are generally two sides of the same coin, especially in the case of polyphenols which can generate hormetic responses (Calabrese et al. 2010). Shikimate has also been implicated in an increased risk of gastric and esophageal cancer found after the consumption of shikimic acid-rich bracken in animals (Evans and Osman 1974; Wilson et al. 1998). Shikimic acid has been proposed to be a carcinogen-promoting agent (Jones et al. 1983). A recent study also found that shikimate can stimulate proliferation of a human breast cancer cell line (MCF-7) via activation of NF-κB signalling (Ma and Ning 2019). The novel mechanism of action of glyphosate on the gut microbiome we describe in the study presented here might be of relevance in the debate on glyphosate’s ability to act as a carcinogen (Guyton et al. 2015). Increased levels of shikimic acid caused by glyphosate inhibition of EPSPS may result in shikimate acting as a potential cancer promoting agent not only in the gut, but also at distant internal organ locations if absorbed. However, further studies should be done using, for example, germ-free mice to understand if the tumour promoting properties, which have been proposed in some studies with glyphosate (George et al. 2010) can be mediated by an action on the gut microbiome.

Shotgun metagenomics revealed that glyphosate changed the composition of the rat gut microbiome (Figure 3). Although large intragroup variations limit the reliability of the results for a large number of statistically significant differences, some taxonomic groups were affected consistently, and in a dose dependent manner. *Eggerthella spp.* had their abundance increased by both glyphosate and MON 52776. This may be due to these bacteria being able to metabolise glyphosate and use it as a source of energy. Although the existence of bacteria metabolising glyphosate in the gut remains unknown, several studies have reported their existence in environmental settings (Grube et al. 2019). It could also be hypothesised that *Eggerthella* spp are more resistant than other bacteria to glyphosate growth inhibiting effects. *Eggerthella spp.* are found in the human gut microbiome (prevalence ∼ 5.5% in Nayfach et al. 2019). Some *Eggerthella spp.* such as *Eggerthella lenta* are known pathogens commonly associated with infections of the gastrointestinal tract (Gardiner et al. 2015). The metabolic capabilities of *Eggerthella spp.* has consequences in humans. This taxa is known to affect the efficacy of drugs used to treat cardiac conditions (Haiser et al. 2013). In addition, changes in *Eggerthella spp.* abundance in the gut have been shown to influence the levels of phenol sulphate, cinnamoylglycine and 5-androstane-3,17-diol in the humsn blood metabolome (Visconti et al. 2019). Interestingly, cinnamoylglycine and glyphosate are both glycine derivatives, which could potentially indicate that they share a common metabolic fate in *Eggerthella spp*. Further studies will be necessary to clarify the dynamic of the gut microbiome compositional alterations induced by glyphosate.

One of the most studied aspect of glyphosate toxicology is its oxidative stress inducing properties. Glyphosate is known to affect mitochondria in mammalian cells, generating oxidative stress (Burchfield et al. 2019; El-Shenawy 2009). Caecal metabolomics in our study showed that glyphosate affected gamma-glutamyl amino acid metabolism in the microbiome (Figure 2). Both gamma-glutamyl amino acid and the dipeptide cysteinylglycine, which are derived from the breakdown of glutathione, were increased. The shotgun metagenomics revealed that the gene encoding the enzyme gamma-glutamyl transpeptidase was decreased from the lowest dose tested (Figure 4B). This can be potentially connected to glyphosate inhibition of EPSPS, since studies in plants have shown that glyphosate-dependent inhibition of shikimic acid metabolism via EPSPS affects cellular redox homeostasis (Vivancos et al. 2011). However, further experiments including more functional readouts such as metatranscriptomics or metaproteomics will be necessary to understand the relevance to health of these findings.

It is also not clear if the gut microbiome is mediating the effects of MON 52776 and glyphosate on the liver. The gut microbiome and liver can regulate each other by a reciprocal communication involving bile acids, antimicrobial molecules and dietary metabolites (Tripathi et al. 2018). Our observation that the formulated product MON 52776 caused adverse effects on liver (Figure 1), is consistent with our previous findings that chronic exposure to an ultra-low dose of Roundup resulted in non-alcoholic fatty liver disease (Mesnage et al. 2015; Mesnage et al. 2017). Interestingly, the strain of *Eggerthella spp*. found to be increased in abundance by glyphosate in our study has been reported to be associated with liver cirrhosis in human populations (Nayfach et al. 2019). However, a direct toxic effect of MON 52776 and glyphosate on liver requires further investigation to understand if glyphosate-mediated increases in the amount of shikimic acid in the caecum microbiome could have an influence on rat health. Longer-term experiments with larger groups of animals will be needed to ascertain if deleterious effects can arise on liver and kidney function. These future experiments could include a exposure at a prenatal period of development in order to ascertain life-long effects (Landrigan and Belpoggi 2018).

Gut microbiome metagenomics and metabolomics can be confounded by a large number of factors, which remain largely unidentified (McLaren et al. 2019). The identification of taxonomic differences was limited by different variables such as the relatively low statistical power provided by the use of 12 animals per group, the incompleteness of the taxonomic classification in gene catalogues, as well as intrinsic factors such as the zero-inflation of metagenomic gene count data (Knight et al. 2012). In addition, different classifiers have been shown to provide different results and there is no gold standard method by which to analyse shotgun metagenomics datasets (Ye et al. 2019). Even the type of instrumentation employed can play a role, with NovaSeq sequencers detecting more DNA sequence diversity within samples than MiSeq sequencers at the exact same sequencing depth (Singer et al. 2019). This could be amplified in our study by the housing of three rats per cage, since rats can exchange faecal microorganisms and metabolites by coprophagy (Suckow et al. 2005).

Many studies have reported that commercially formulated pesticide products are more toxic than their active ingredient. This has been shown multiple times for glyphosate-based herbicides (Adam et al. 1997; Manservisi et al. 2019; Mesnage et al. 2013), and also for other classes of pesticides such as insecticides or fungicides (Mesnage et al. 2014). This difference in toxic effects is generally due to the toxicity of surfactants and other compounds that constitute the adjuvant mixture in commercial pesticide formulations (Mesnage et al. 2019), although it cannot be excluded that these surfactants can also potentiate glyphosate penetration in tissues (Anadón et al. 2009). Few studies have examined the toxicity of compounds used as pesticide co-formulants on the gut microbiome, with the only comprehensive study published so far suggesting that compounds having emulsifying properties can drive intestinal inflammation by affecting the gut mucosa (Chassaing et al. 2017). Our study suggests that the adjuvant mixture present in MON 52776 had limited effects on the caecum metabolome in comparison to glyphosate, which was the main ingredient responsible for the metabolic changes observed in our study. There were nonetheless taxonomic differences with *Shinella zoogleoides* found to be increased by exposure to MON 52776 but not glyphosate. The potential roles of *Shinella spp.* in the gut microbiome are still elusive, although it is notable that some *Shinella spp.* have been isolated from various environmental samples, such as active sludge, and are known to degrade environmental pollutants including chlorothalonil (Liang et al. 2011) and the alkaloid nicotine (Qiu et al. 2016). We hypothesise that the increase in *Shinella* spp. caused by MON 52776 could affect alkaloid levels in the gut as suggested by the large decrease in solanidine levels, which was only detected in the MON 52776 treated group. We also found that different classes of surfactants used in glyphosate formulated products can have different toxicity profiles on bacteria (Figure 5), suggesting that results with one formulation should not be generalised to all other GBH products.

In conclusion, our study demonstrates the power of using multi-omics molecular profiling to reveal changes in the gut microbiome following exposure to chemical pollutants that would otherwise be missed using more standard, less comprehensive analytical methods. Employing this approach allowed us to identify the first biomarker of glyphosate effects on the rat gut microbiota, namely a marked increase in shikimate and 3-dehydroshikimate reflective of inhibition of the EPSPS enzyme of the shikimate pathway. In addition, we found increased levels of γ-glutamylglutamine, cysteinylglycine and valylglycine suggestive of a response to oxidative stress. Furthermore, our results showed distinct changes in the profile of microbiota with glyphosate causing increased levels of *Eggerthella spp*. and *Homeothermacea spp* whilst MON 52276, increased the relative abundance of *Shinella zoogleoides*. These shifts in bacterial species, once confirmed by further studies, could also act as additional biomarkers of glyphosate and Roundup exposure. Although more studies are needed to understand the health implications of glyphosate inhibition of the shikimate pathway in the gut microbiome, our findings can be used in environmental epidemiological studies to understand if glyphosate can have biological effects in human populations.

## Supporting information

Supplementary Material 1. Feed analysis to identify possible contaminants.Supplementary Material 2. Shotgun metagenomics analysis at the phylum level.

Supplementary Material 2. Shotgun metagenomics analysis at the phylum level.

## Acknowledgements

We thank Dr. Nicola Segata and Dr. Francesco Asnicar for assistance in metagenomics data analysis using their pre-processing package and useful discussions on metagenomics analytical pipelines. This work was funded by the Sustainable Food Alliance (USA) and in part by the Sheepdrove Trust (UK).

## Supplementary Materials

**Supplementary Material 1.** Feed analysis to identify possible contaminants.

**Supplementary Material 2.** Shotgun metagenomics analysis at the phylum level.

