## Supplementary Material 1. Feed analysis to identify possible contaminants.Supplementary Material 2. Shotgun metagenomics analysis at the phylum level. for "Shotgun metagenomics and metabolomics reveal glyphosate alters the gut microbiome of Sprague-Dawley rats by inhibiting the shikimate pathway"

SAFE
COMPLETE CARE COMPETENCE
www.safe-diets.com

### Rats, Mice & Hamsters maintenance diet

### SAFE A04

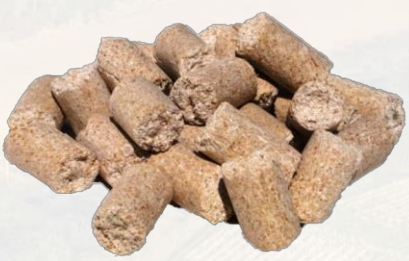


#### Definition

Complete maintenance diet for rats, mice & hamsters.

#### Product Purpose

Rodent diet for adult and maintenance animals.

To be used within the context of experimental protocols. Does not contain Alfalfa.

**Distribution period:** from weaning and to adult rodents.

**Daily consumption:** rats 18 to 25 g, mice 3 to 6 g, hamsters 8 to 12g.

**Distribution method:** ad libitum or rationed according to experimental protocols.

#### Product Presentation

15 mm diameter pellet. Can be modified on demand.

| Diet | Packaging | Control sheet | Irradiation dose | Animals status | Product code |
|---|---|---|---|---|---|
| **SAFE A04** | Paper bag 10 kg | No | None | Conventionnal | U8220G10R |
| **SAFE A04-10** | Double paper bag 10 kg | No | >10 kiloGrays | Heteroxenic | U8221G10R |
| **SAFE A04 SP-10** | Paper bag in hermetic plastic pouch 10 kg | No | >10 kiloGrays | Heteroxenic | U8996G10R |
| **SAFE A04 SP-25** | Paper bag in hermetic plastic Pouch 10 kg | No | >25 kiloGrays | Heteroxenic | U8993G10R |
| **SAFE R04-10** | Paper bag vacuum packed box of 1x10 kg | No | >10 kiloGrays | Heteroxenic | U8231G10R |
| **SAFE R04-25** | Paper bag vacuum packed box of 1x10 kg | No | >25 kiloGrays | Heteroxenic | U8232G10R |
| **SAFE R04-40** 10x1 kg | Double vacuum packed box of 10x1 kg | No | >40 kiloGrays | Axenic and Gnotoxenic | U8233G10R |
| **SAFE A04C** | Double paper bag 10 kg | Yes | None | Conventionnal | U8224G10R |
| **SAFE A04C-10** | Double paper bag 10 kg | Yes | >10 kiloGrays | Heteroxenic | U8225G10R |

All diets are available with custom packaging, irradiated and with a complete analysis on demand. They are available powdered on demand.

Updated October 05th 2017



SAFE
COMPLETE CARE COMPETENCE
www.safe-diets.com

### Rats, Mice & Hamsters maintenance diet

### SAFE A04

#### Nutritional Composition/kg

| AMINO ACIDS | | TOTAL * |
|---|---|---|
| Arginine | mg | 9 000 |
| Cystine | mg | 2 500 |
| Lysine | mg | 7 200 |
| Méthionine | mg | 2 800 |
| Tryptophane | mg | 1 900 |
| Glycine | mg | 8 100 |

| FATTY ACIDS | | TOTAL * |
|---|---|---|
| Palmitic acid | mg | 5 900 |
| Plamitoleic acid | mg | 150 |
| Stearic acid | mg | 600 |
| Oleic acid | mg | 4 800 |
| Linoleic acid | mg | 15 000 |
| Linolenic acid | mg | 1 200 |

| MINERALS | | TOTAL * |
|---|---|---|
| P | mg | 5 500 |
| Ca | mg | 7 300 |
| Na | mg | 2 500 |
| K | mg | 6 000 |
| Mg | mg | 1 600 |
| Mn | mg | 70 |
| Fe | mg | 270 |
| Cu | mg | 16 |
| Zn | mg | 55 |
| Cl | mg | 4 000 |

| VITAMINS | | TOTAL * |
|---|---|---|
| Vitamin A | UI | 7 500 |
| Vitamin D3 | UI | 1 000 |
| Vitamin B1 | mg | 5 |
| Vitamin B2 | mg | 6,5 |
| Vitamin B5 | mg | 10 |
| Vitamin B6 | mg | 3 |
| Vitamin B12 | mg | 0,01 |
| Vitamin E | UI | 30 |
| Vitamin K3 | mg | 2,5 |
| Niacin | mg | 70 |
| Folic Ac. | mg | 0,35 |
| Biotin | mg | 0,08 |
| Choline | mg | 1 600 |

#### Pellets Technology

| | | Mean* |
|---|---|---|
| Diameter | mm | 16,43 |
| Resistance to crushing | kgf/cm² | 22, 7 |
| Resistance to abrasing | % | 97,3 |
| Specific mass | g/l | 645 |
| Average pellet weight | g | 5,319 |
| Average pellet length | mm | 22,64 |

#### Composition

Barley, wheat, maize, soybean meal, wheat bran, hydrolyzed fish proteins, dicalcium phosphate, pre-mixture of minerals, calcium carbonate, pre-mixture of vitamins.

#### Centesimal Composition in %

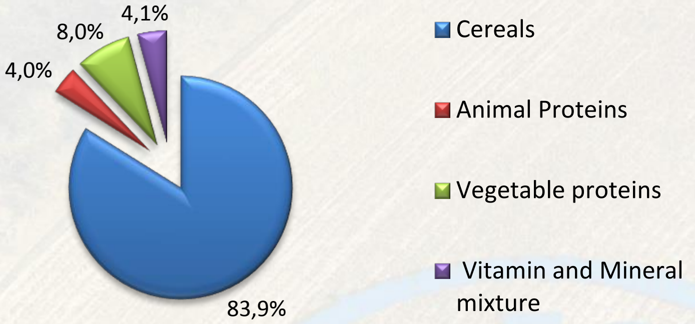


#### Nutritional Composition in %

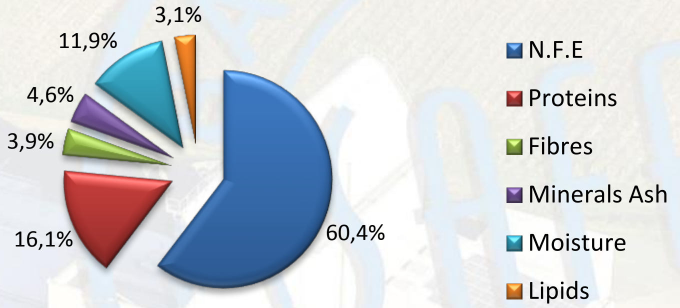


| Energy value** | Kcal/kg | Mj/kg | % Proteins | % Lipids | % Carbohydrates |
|---|---|---|---|---|---|
| Atwater | 3 339 | 13,97 | 19,3 | 8,4 | 72,4 |
| ME | 3 145 | 13,17 | - | - | - |

** Energy calcul information:
http://www.safe-diets.com/en/services-r-and-d/diet-energy/

| Nitrogen free extract | | |
|---|---|---|
| - of which starch | (%) | 43.5 |
| - of which total sugars | (%) | 3.2 |

* Values are given as an indication only. They are calculated averages of product raw values. They are indicative and have no contractual value. They are subject to variations related to production conditions storage and analytical methods. An analysis is performed on request.

N.F.E.: Nitrogen-free extract, calculated value.

Updated October 05th 2017

### Rapport d'analyse

| Echantillon n° | 370-2017-00194930 | Date 24/08/2017 | Page 1/3 |
|---|---|---|---|
| Rapport d'analyse n° | AR-17-AA-186115-01 / 370-2017-00194930 | | |

SAFE

A l'attention de **Monsieur Renaud BARRAL**
Route de Saint Bris
89290 AUGY
FRANCE

**Copie à :** Monsieur Martel

**Coordinateur technique de votre dossier :** Marion Greloux

| | | | |
|---|---|---|---|
| **Notre référence :** | 370-2017-00194930/ AR-17-AA-186115-01 | **Type :** | EX |
| **Référence client :** | **U8220 V257, CD011344** | | |
| **Description de l'échantillon :** | 80% céréales | | |
| **Conditionnement :** | NonCommercial : 672g | | |
| **Votre date de commande :** | 10/08/2017 | **Votre référence commande :** | 10/07/2017 / (EOL) 518-624026 |
| **Date de réception :** | 17/08/2017 12:40:00 | **Date de mise en analyse :** | 17/08/2017 |
| **Prélèvement/Transport :** | dpd | | |
| **Analyses demandées :** | AAU : Formule OC/OP/PYR/naled/temephos/PCB<br>SF00B : Glyphosate, glufosinate, AMPA (alimentation) | | |
| **DLC/DLUO** | / | **N° de lot** | 17207 |
| **Marque** | envoi E8797 | **Code emballeur** | RB |

| | | | Résultats (incertitude) |
|---|---|---|---|
| **SFW9Z** | **SF** | **Analytical run test GC** | |
| | | Analytical run | FAIT |

| Pesticides | | | Résultats (incertitude) |
|---|---|---|---|
| **SF00B** | **SF** | **Glyphosate, glufosinate, AMPA (alimentation) Méthode : Méthode interne, LC/MS/MS** | |
| | (a) | Pesticides recherchés | <LOQ |
| **SFVNS** | **SF** | **PCB avec Screening GC/MS Méthode : § 64 LFGB L 00.00-38 1-4** | |
| | (a) | PCB 101 | <0.005 mg/kg |
| | (a) | PCB 118 | <0.005 mg/kg |
| | (a) | PCB 138 | <0.005 mg/kg |
| | (a) | PCB 153 | <0.005 mg/kg |
| | (a) | PCB 180 | <0.005 mg/kg |
| | (a) | PCB 28 | <0.005 mg/kg |
| | (a) | PCB 52 | <0.005 mg/kg |
| **SFLA0** | **SF** | **Screening pesticides (GC/MS) Méthode : §64 LFGB L00.00-34, mod.** | |
| | (a) | Pesticides recherchés | <LOQ |
| **SFLD0** | **SF** | **Screening pesticides (LC/MS/MS) Méthode : LFGB L 00.00-113** | |
| | (a) | Butoxyde de Pipéronyle (PBO) | 0.034 (± 0.017) mg/kg |
| | (a) | Autres pesticides recherchés | <LOQ |
| **SFCEB** | **SF** | **Naled / SF0XA Méthode : Méthode interne, GC/MS** | |
| | (a) | Naled | <0.05 mg/kg |

**CONCLUSION (Non couverte par l'accréditation)**

Aux limites de quantification des méthodes mises en oeuvre, aucun des paramètres suivants n'a été observé :
somme des 6 PCBs < 0.005 mg/kg
somme de l'Heptachlore et de l'Heptachlorepoxide- cis et -trans < 0.005 mg/kg"

#### Liste des molécules recherchées et non détectées (* = limite de quantification)

**SF00B SF Glyphosate, glufosinate, AMPA (alimentation)** (LOQ* mg/kg)

(a) Acide aminométhylphosphonique (AMPA) (.01 ) (a) Glufosinate (.01 ) (a) Glyphosate (.01 ) eurofins | analytics

**Echantillon n°** **370-2017-00194930** **Date 24/08/2017** **Page 2/3**
**Rapport d'analyse n°** **AR-17-AA-186115-01 / 370-2017-00194930**

**SFLA0 SF Screening pesticides (GC/MS)** (LOQ* mg/kg)

| | | | | | |
|---|---|---|---|---|---|
| (a) 2,4,5-T-Méthylester (0.01) | (a) 2,4-D-Méthyl Ester (0.01) | (a) 4,4-Dibromobenzophénone (0.01) | (a) Acetochlor (0.05) | (a) Aclonifen (0.02) | (a) Acrinathrine (0.01) |
| (a) Alachlore (0.1) | (a) Aldrine (0.005) | (a) Alléthrine (Dépalléthrine) (0.01) | (a) Amidithion (0.01) | (a) Atrazine (0.1) | (a) Azaconazole (0.05) |
| (a) Azinphos-ethyl (0.01) | (a) Azinphos-methyl (0.01) | (a) Azoxystrobine (0.01) | (a) Benfluraline (0.005) | (a) Benoxacor (0.01) | (a) Benzoylprop-ethyl (0.05) |
| (a) Béta-endosulfan (0.005) | (a) Bifénox (0.01) | (a) Bifenthrine (0.005) | (a) Binapacryl (0.02) | (a) Bitertanol (0.1) | (a) Boscalide (0.05) |
| (a) Bromocyclen (0.01) | (a) Bromofenvinphos (0.01) | (a) Bromophos-ethyl (0.005) | (a) Bromophos-méthyl (0.005) | (a) Bromopropylate (0.01) | (a) Buprofezine (0.05) |
| (a) Butachlore (0.01) | (a) Butamifos (0.005) | (a) Butraline (0.01) | (a) Cadusaphos (0.01) | (a) Captafol (0.01) | (a) Captane (0.01) |
| (a) Carbophenothion (0.005) | (a) Carbophénothion-méthyl (0.01) | (a) Carfentrazone-ethyl (0.01) | (a) Chinomethionate (0.01) | (a) Chlorbenside (0.005) | (a) Chlordane-cis (0.005) |
| (a) Chlordane-gamma (=bêta=trans) (0.005) | (a) Chlordécone (0.01) | (a) Chlorethoxyfos (0.005) | (a) Chlorfenapyr (0.005) | (a) Chlorfenprop-méthyl (0.01) | (a) Chlorfenson (0.005) |
| (a) Chlorfenvinphos (0.01) | (a) Chloridazon (Pyrazon) (0.1) | (a) Chlorméphos (0.005) | (a) Chlorobenzilate (0.005) | (a) Chloroneb (0.1) | (a) Chloropropylate (0.005) |
| (a) Chlorothalonil (0.005) | (a) Chlorpyrifos (-ethyl) (0.005) | (a) Chlorpyrifos-méthyl (0.005) | (a) Chlorthal diméthyle (0.005) | (a) Chlorthion (0.01) | (a) Chlorthiophos (0.005) |
| (a) Chlozolinate (0.01) | (a) Cinidon-éthyle (0.025) | (a) Clodinafop-propargyl (0.03) | (a) Coumaphos (0.005) | (a) Crotoxyphos (0.01) | (a) Cyanofenphos (0.01) |
| (a) Cyanophos (0.01) | (a) Cyfluthrine (0.01) | (a) Cyhalothrine (0.01) | (a) Cyperméthrine (0.005) | (a) Cyphenothrine (0.01) | (a) Cyproconazole (0.05) |
| (a) DDD, o,p (0.005) | (a) DDE, o,p' (0.005) | (a) DDT (p,p'-DDT+o,p'-DDT+p,p'-DDE+p,p'-TDE) (0.01) | (a) Deltamethrine (0.005) | (a) Dialifos (0.01) | (a) Diallat (0.01) |
| (a) Diazinon (0.01) | (a) Dicapthon (0.01) | (a) Dichlobénil (0.05) | (a) Dichlofenthion (0.01) | (a) Dichlofluanide (0.005) | (a) Dichloran (0.005) |
| (a) Dichlorvos (0.025) | (a) Diclofop-méthyl (0.02) | (a) Dicofol (0.005) | (a) Dicofol, o,p- (0.005) | (a) Dicrotophos (0.01) | (a) Dieldrine (0.005) |
| (a) Dienochlor (0.01) | (a) Difénoconazole (0.02) | (a) Diflufenican (0.01) | (a) Dimefox (0.05) | (a) Dimethachlor (0.01) | (a) Dimethipine (0.01) |
| (a) Diméthoate (0.02) | (a) Dimethomorphe (0.05) | (a) Diniconazole (0.01) | (a) Dinitramine (0.01) | (a) Dinobuton (0.05) | (a) Disulfoton (0.05) |
| (a) Disulfoton sulfone (0.02) | (a) Ditalimphos (0.005) | (a) Edifenphos (0.01) | (a) Endosulfan alpha (0.005) | (a) Endosulfan sulfate (0.005) | (a) Endrine (0.005) |
| (a) EPN (0.01) | (a) Epoxyconazole (0.04) | (a) Etaconazole (0.02) | (a) Ethalfluraline (0.01) | (a) Ethion (0.005) | (a) Ethiprol (0.02) |
| (a) Ethofumesate (0.2) | (a) Ethoprophos (0.005) | (a) Ethyl parathion (0.005) | (a) Etridiazole (0.005) | (a) Etrimphos (0.005) | (a) Famophos (0.01) |
| (a) Famoxadone (0.01) | (a) Fenamidone (0.02) | (a) Fenamiphos (0.02) | (a) Fénarimol (0.05) | (a) Fenbuconazole (0.1) | (a) Fenchlorazol (0.01) |
| (a) Fenchlorphos (0.005) | (a) Fenfluthrine (0.01) | (a) Fenhexamid (0.05) | (a) Fénitrothion (0.005) | (a) Fenoxaprop-éthyle (0.05) | (a) Fenpiclonil (0.05) |
| (a) Fenpropathrine (0.005) | (a) Fenpropimorphe (0.1) | (a) Fenson (0.01) | (a) Fensulfothion (0.01) | (a) Fenvalerate (RR-/SS-Isomere) (0.005) | (a) Fenvalerate (RS-/SR-Isomere) (0.005) |
| (a) Fipronil (0.004) | (a) Fipronil desulfinyl (0.004) | (a) Fipronil sulfite (0.005) | (a) Fipronil sulfon (0.005) | (a) Flamprop-isopropyle (0.01) | (a) Flamprop-méthyl (0.01) |
| (a) Flonicamide (0.05) | (a) Fluazifop-butyl (0.05) | (a) Fluaziname (0.01) | (a) Fluchloraline (0.005) | (a) Flucythrinate (0.01) | (a) Flufenoxuron (0.01) |
| (a) Fluméthrine (0.01) | (a) Flumetraline (0.005) | (a) Fluopicolid (0.01) | (a) Fluorodifen (0.005) | (a) Fluotrimazole (0.01) | (a) Fluquinconazole (0.01) |
| (a) Flurenol-Butyl (0.02) | (a) Flurochloridone (0.05) | (a) Flurtamone (0.01) | (a) Flusilazole (0.1) | (a) Folpel (Folpet) (0.01) | (a) Fonofos (0.005) |
| (a) Formothion (0.01) | (a) Genite (0.01) | (a) Halfenprox (0.01) | (a) Haloxyfop-Ethoxyéthylé (0.01) | (a) Haloxyfop-methyl (0.05) | (a) HCH Alpha (0.005) |
| (a) HCH Béta (0.005) | (a) HCH Delta (0.005) | (a) HCH, gamma - Lindane (0.005) | (a) HCH-epsilon (0.005) | (a) Heptachlore (0.005) | (a) Heptachlore époxide cis (0.005) |
| (a) Heptachlore époxide trans (0.005) | (a) Hepténophos (0.01) | (a) Hexachlorobenzène (HCB) (0.005) | (a) Hexaconazole (0.015) | (a) IBP (Iprobenfos) (0.01) | (a) Indanofan (0.02) |
| (a) Indoxacarbe (0.005) | (a) Iodofenphos (0.005) | (a) Ioxynil-Octanoate (0.01) | (a) Iprodione (0.01) | (a) Isazophos (0.01) | (a) Isobenzane (0.005) |
| (a) Isocarbofos (0.01) | (a) Isodrine (0.005) | (a) Isofenphos (0.005) | (a) Isofenphos-Méthyl (0.005) | (a) Isomethiozin (0.01) | (a) Isopropalin (0.005) |
| (a) Isoxadifen-éthyle (0.01) | (a) Kétoendrin-delta (0.005) | (a) Kresoxime-méthyl (0.005) | (a) Lactofen (0.01) | (a) Lambda cyhalothrine (0.01) | (a) Leptophos (0.01) |
| (a) Lufénuron (0.012) | (a) Malaoxon (degradation Malathion) (0.01) | (a) Malathion (0.005) | (a) Mecarbam (0.01) | (a) Mephosfolan (0.01) | (a) Merphos (0.01) |
| (a) Métazachlore (0.01) | (a) Méthacrifos (0.05) | (a) Méthidathion (0.012) | (a) Méthoxychlore (0.005) | (a) Métolachlore (0.05) | (a) Metrafenone (0.05) |
| (a) Métribuzine (0.01) | (a) Mévinphos (0.01) | (a) Mirex (0.005) | (a) Molinate (0.1) | (a) Myclobutanile (0.005) | (a) Nitralin (0.01) |
| (a) Nitrapyrine (0.02) | (a) Nitrofene (0.01) | (a) Nitrothal-isopropyle (0.01) | (a) Norflurazon (0.05) | (a) Nuarimol (0.01) | (a) Ométhoate (0.1) |
| (a) Oxadiazon (0.005) | (a) Oxychlordane (0.01) | (a) Oxydéméton methyl (0.02) | (a) Oxyfluorfene (0.01) | (a) Paclobutrazole (0.04) | (a) Paraoxon (0.02) |
| (a) Paraoxon-méthyl (0.02) | (a) Parathion-méthyl (0.005) | (a) Penconazole (0.02) | (a) Pendiméthaline (0.005) | (a) Pentachloraniline (0.01) | (a) Pentachloroanisole (PCA) (0.005) |
| (a) Pentachlorobenzène (0.005) | (a) Pentachlorothioanisole (0.01) | (a) Perméthrine (0.02) | (a) Perthane (0.2) | (a) Phenkapton (0.01) | (a) Phénothrine (0.01) |
| (a) Phenthoate (0.01) | (a) Phosalone (0.01) | (a) Phosfolane (0.01) | (a) Phosmet (0.02) | (a) Picolinafene (0.01) | (a) Picoxystrobin (0.02) |
| (a) Piperophos (0.01) | (a) Pirimiphos-ethyl (0.01) | (a) Pirimiphos-méthyl (0.01) | (a) Pirimitate (0.05) | (a) Plifenate (0.01) | (a) Pralléthrine (0.01) |
| (a) Procymidone (0.015) | (a) Profenofos (0.005) | (a) Profluraline (0.005) | (a) Propachlore (0.01) | (a) Propanile (0.01) | (a) Propazine (0.05) |
| (a) Propétamphos (0.005) | (a) Propiconazole (0.01) | (a) Propyzamide (0.01) | (a) Prothiophos (0.005) | (a) Prothoate (0.01) | (a) Pyraclofos (0.01) |
| (a) Pyraflufen-éthyl (0.02) | (a) Pyrazophos (0.01) | (a) Pyréthrines (total) (0.02) | (a) Pyridabène (0.005) | (a) Pyridaphenthion (0.01) | (a) Pyrifenox (0.01) |
| (a) Quinalphos (0.005) | (a) Quinoxyfen (0.015) | (a) Quintozène (0.005) | (a) Quizalofop ethyle (0.02) | (a) Resmethrin (0.1) | (a) S 421 (0.01) |
| (a) Spiromesifene (0.01) | (a) Sulfotep (0.025) | (a) Sulprofos (0.01) | (a) Swep (0.01) | (a) Tau-fluvalinate (0.005) | (a) Tebupirimfos (0.005) |
| (a) Tecnazéne (0.005) | (a) Téfluthrine (0.005) | (a) Temephos (0.01) | (a) Terbufos (0.005) | (a) Tetrachlorvinphos (0.005) | (a) Tetraconazole (0.025) |
| (a) Tétradifon (0.005) | (a) Tétramethrine (0.01) | (a) Tétrasul (0.01) | (a) Tolclofos-méthyl (0.01) | (a) Tolylfluanide (0.01) | (a) Toxaphène Parlar N°26 (0.01) |
| (a) Toxaphène Parlar N°50 (0.01) | (a) Toxaphène Parlar N°62 (0.01) | (a) Transfluthrin (0.01) | (a) Triadimefone (0.01) | (a) Triadimenole (0.1) | (a) Triallate (0.01) |
| (a) Triamiphos (0.025) | (a) Triazophos (0.02) | (a) Tribufos (0.01) | (a) Trichloronate (0.005) | (a) Tridiphane (0.05) | (a) Trifloxystrobine (0.01) |
| (a) Trifluraline (0.005) | (a) Vamidothion (0.05) | (a) Vinclozoline (0.005) | | | |

**SFLD0 SF Screening pesticides (LC/MS/MS)** (LOQ* mg/kg)

| | | | | | |
|---|---|---|---|---|---|
| (a) 2,4'-Formoxylidid (métabolite de l'Amitraze) (0.01) | (a) 3-Hydroxycarbofurane (0.005) | (a) 5-Hydroxy-Thiabendazol (0.01) | (a) 6-Chloro-3-phenyl pyridazin-4-ol (0.005) | (a) Abamectine (0.01) | (a) Acéphate (0.005) |
| (a) Acétamipride (0.005) | (a) Acetochlor (0.02) | (a) Alachlore (0.01) | (a) Aldicarb sulfone (0.01) | (a) Aldicarb sulfoxyde (0.005) | (a) Aldicarbe (0.005) |
| (a) Ametoctradin (0.01) | (a) Amétryne (0.005) | (a) Amidosulfuron (0.01) | (a) Aminocarbe (0.01) | (a) Amitraze (0.01) | (a) Ancymidol (0.01) |
| (a) Atrazine (0.005) | (a) Azaconazole (0.01) | (a) Azaméthiphos (0.01) | (a) Aziprotryn (0.01) | (a) Azoxystrobine (0.005) | (a) Bénalaxyl (0.01) |
| (a) Bendiocarbe (0.005) | (a) Benfuracarbe (0.005) | (a) Benodanil (0.005) | (a) Bénomyl (0.005) | (a) Bensulfuron méthyle (0.005) | (a) Benthiavalicarb-isopropyl (0.005) |
| (a) Bitertanol (0.01) | (a) Boscalide (0.005) | (a) Bromacile (0.01) | (a) Bupirimate (0.01) | (a) Buprofezine (0.005) | (a) Butachlore (0.02) |
| (a) Butocarboxim (0.005) | (a) Butocarboxim sulfoxide (0.005) | (a) Butoxycarboxim (0.005) | (a) Buturon (0.005) | (a) Cadusaphos (0.01) | (a) Carbaryl (0.005) |
| (a) Carbendazime (0.005) | (a) Carbofuran (0.005) | (a) Carbosulfan (0.005) | (a) Carboxine (0.005) | (a) Chlorantraniliprole (0.01) | (a) Chlorbromuron (0.005) |
| (a) Chlorfluazuron (0.01) | (a) Chloridazon (Pyrazon) (0.005) | (a) Chloroxuron (0.01) | (a) Chlorpropham (0.02) | (a) Chlorsulfuron (0.005) | (a) Chlortoluron (0.005) |
| (a) Cinidon-éthyle (0.01) | (a) Cinosulfuron (0.005) | (a) Clethodim (0.01) | (a) Clodinafop-propargyl (0.01) | (a) Clofentézine (0.005) | (a) Clomazone (0.005) |
| (a) Clothianidin (0.01) | (a) Cyanazine (0.01) | (a) Cyazofamide (0.02) | (a) Cymoxanil (0.1) | (a) Cyproconazole (0.005) | (a) Cyprodinile (0.01) |
| (a) Cyprofuram (0.03) | (a) Cyromazine (0.02) | (a) DEET Diethyltoluamide (0.01) | (a) Demeton (0.01) | (a) Demeton-S-méthyl (0.005) | (a) Demeton-S-methyl-sulfone (0.005) |
| (a) Desethyl-atrazine (0.005) | (a) Desethyl-Simazine (Désisopropylatrazine) (0.01) | (a) Deséthyl-terbutylazine (0.005) | (a) Desmedipham (0.005) | (a) Desmetryne (0.005) | (a) Diazinon (0.01) |
| (a) Dichlorvos (0.005) | (a) Diclobutrazole (0.01) | (a) Diethofencarbe (0.005) | (a) Difénoconazole (0.005) | (a) Difénoxuron (0.01) | (a) Diflubenzuron (0.02) |
| (a) Diflufenican (0.01) | (a) Dimefox (0.05) | (a) Dimefuron (0.01) | (a) Diméthénamide (0.005) | (a) Diméthoate (0.005) | (a) Dimethomorphe (0.01) |

**Eurofins Analytics France (Nantes)**
Rue Pierre Adolphe Bobierre
BP 42301
F-44323 Nantes Cedex 3
**FRANCE**

Tél. +33 2 51 83 43 40

www.eurofins.fr

SAS au capital de 3 256 700 €
RCS NANTES 423 190 891
SIRET 423 190 891 00022
APE 743 B

eurofins | analytics

### Rapport d'analyse

**Echantillon n°** **370-2017-00194930** **Date 24/08/2017** 

**Rapport d'analyse n°** **AR-17-AA-186115-01 / 370-2017-00194930**

**SFLD0 SF Screening pesticides (LC/MS/MS)** (LOQ* mg/kg)

| | | | | | |
|---|---|---|---|---|---|
| (a) Dimetilan (0.005) | (a) Dimoxystrobine (0.005) | (a) Dinotefuran (0.05) | (a) Disulfoton (0.005) | (a) Disulfoton sulfone (0.01) | (a) Disulfoton sulfoxyde (0.01) |
| (a) Diuron (0.005) | (a) Emamectine (Somme) (0.01) | (a) Epoxyconazole (0.005) | (a) Ethiofencarbe (0.005) | (a) Ethiofencarb-sulfone (0.005) | (a) Ethiofencarb-sulfoxyde (0.005) |
| (a) Ethiprol (0.01) | (a) Ethofumésat-2-keto (0.05) | (a) Ethofumesate (0.01) | (a) Ethoprophos (0.005) | (a) Etofenprox (0.005) | (a) Etoxazole (0.01) |
| (a) Famoxadone (0.01) | (a) Fenamidone (0.01) | (a) Fenamiphos (0.01) | (a) Fenamiphos-sulfone (0.01) | (a) Fenamiphos-sulfoxyde (0.01) | (a) Fénarimol (0.005) |
| (a) Fénazaquine (0.005) | (a) Fenbuconazole (0.005) | (a) Fenhexamid (0.005) | (a) Fenobucarb (0.01) | (a) Fenoxaprop-éthyle (0.01) | (a) Fenoxycarbe (0.005) |
| (a) Fenpiclonil (0.01) | (a) Fenpropidin (0.01) | (a) Fenpropimorphe (0.005) | (a) Fenpyroximate (0.01) | (a) Fensulfothion (0.01) | (a) Fensulfothione Sulfone (0.01) |
| (a) Fensulfothion-PO-sulfon (0.01) | (a) Fensulfothion-PO-sulfoxide (0.01) | (a) Fenthion (0.01) | (a) Fenthion-oxone (0.01) | (a) Fenthion-PO-sulfoxid (0.01) | (a) Fenthion-PS-Sulfoxid (0.01) |
| (a) Fention-PO-sulfon (0.01) | (a) Fention-PS-sulfon (0.01) | (a) Fenuron (0.005) | (a) Flazasulfuron (0.005) | (a) Flonicamide (0.01) | (a) Florasulam (0.005) |
| (a) Fluazifop-P-butyle (0.005) | (a) Fluazuron (0.02) | (a) Flucycloxuron (0.005) | (a) Fludioxonil (0.01) | (a) Flufenacet (0.01) | (a) Flufenoxuron (0.01) |
| (a) Fluometuron (0.01) | (a) Fluopicolid (0.01) | (a) Flurochloridone (0.01) | (a) Flurprimidol (0.01) | (a) Flusilazole (0.005) | (a) Flutriafol (0.01) |
| (a) FM-6-1 (metabolite du Triflumizole) (0.01) | (a) Formetanate (0.01) | (a) Fosthiazate (0.01) | (a) Fuberidazole (0.005) | (a) Furathiocarb (0.005) | (a) Halofenozide (0.005) |
| (a) Haloxyfop-Ethoxyéthylé (0.005) | (a) Haloxyfop-methyl (0.005) | (a) Hexaconazole (0.01) | (a) Hexaflumuron (0.05) | (a) Hexazinone (0.01) | (a) Hexythiazox (0.005) |
| (a) Imazalile (0.005) | (a) Imibenconazole (0.01) | (a) Imidaclopride (0.005) | (a) Indoxacarbe (0.005) | (a) Iodosulfuron méthyle (0.005) | (a) Iprovalicarbe (0.005) |
| (a) Isoprocarb (0.01) | (a) Isoprothiolane (0.01) | (a) Isoproturon (0.005) | (a) Isoxaben (0.01) | (a) Isoxaflutole (0.005) | (a) Isoxathion (0.01) |
| (a) Lénacile (0.01) | (a) Linuron (0.005) | (a) Lufénuron (0.01) | (a) Malaoxon (degradation Malathion) (0.01) | (a) Malathion (0.01) | (a) Mandipropamide (0.01) |
| (a) Mepanipyrim (0.005) | (a) Metalaxyl (0.005) | (a) Metamitrone (0.005) | (a) Métazachlore (0.005) | (a) Metconazole (0.01) | (a) Methabenzthiazuron (0.005) |
| (a) Méthacrifos (0.01) | (a) Methamidophos (0.005) | (a) Méthidathion (0.01) | (a) Methiocarb sulfone (0.01) | (a) Methiocarb Sulfoxyde (0.01) | (a) Méthiocarbe (0.005) |
| (a) Méthomyl (0.005) | (a) Methoprotryne (0.005) | (a) Methoxyfenozid (0.005) | (a) Metobromuron (0.005) | (a) Métolachlore (0.005) | (a) Metolcarb (0.005) |
| (a) Métoxuron (0.005) | (a) Metrafenone (0.005) | (a) Métribuzine (0.005) | (a) Metsulfuron méthyle (0.005) | (a) Molinate (0.01) | (a) Monocrotophos (0.005) |
| (a) Monolinuron (0.005) | (a) Monuron (0.005) | (a) N-2,4-dimethylphenyl-N-methylformamidine (0.05) | (a) Napropamide (0.01) | (a) Néburon (0.005) | (a) Nicosulfuron (0.01) |
| (a) Novaluron (0.01) | (a) Nuarimol (0.025) | (a) Ofurace (0.005) | (a) Ométhoate (0.01) | (a) Orbencarb (0.005) | (a) Oxadixyl (0.005) |
| (a) Oxamyl (0.005) | (a) Oxamyl-oxime (0.01) | (a) Oxydéméton methyl (0.005) | (a) Paclobutrazole (0.01) | (a) Paraoxon (0.01) | (a) Paraoxon-méthyl (0.01) |
| (a) Penconazole (0.005) | (a) Pencycuron (0.01) | (a) Pendiméthaline (0.005) | (a) Pentanochlor (0.01) | (a) Phenmédiphame (0.005) | (a) Phorate (0.01) |
| (a) Phorate sulfoxyde (0.01) | (a) Phorat-sulfon (0.01) | (a) Phosmet (0.01) | (a) Phosphamidon (0.01) | (a) Phoxime (0.01) | (a) Picoxystrobin (0.005) |
| (a) Pirimicarb, desmethyl-formamido- (0.005) | (a) Pirimicarbe (0.005) | (a) Pirimicarbe, Desmethyl- (0.005) | (a) Primsulfuron methyl (0.005) | (a) Prochloraz (0.005) | (a) Promecarb (0.005) |
| (a) Prométone (0.005) | (a) Prométryne (0.005) | (a) Propamocarbe (0.005) | (a) Propargite (0.01) | (a) Propazine (0.005) | (a) Prophame (0.03) |
| (a) Propiconazole (0.01) | (a) Propoxur (0.005) | (a) Propoxycarbazone (0.01) | (a) Proquinazid (0.005) | (a) Prosulfocarbe (0.01) | (a) Prosulfuron (0.005) |
| (a) Pymétrozine (0.005) | (a) Pyraclostrobine (0.005) | (a) Pyraflufen-éthyl (0.01) | (a) Pyréthrines (total) (0.5) | (a) Pyridate (0.005) | (a) Pyriméthanil (0.005) |
| (a) Pyrimidifen (0.01) | (a) Pyriproxyfen (0.005) | (a) Quizalofop ethyle (0.005) | (a) Rabenzazole (0.005) | (a) Rimsulfuron (0.02) | (a) Rotenone (0.01) |
| (a) Sebuthylazine (0.005) | (a) Sethoxydim (0.01) | (a) Silafluofen (0.05) | (a) Simazine (0.005) | (a) Simeconazole (0.005) | (a) Spinosad (0.005) |
| (a) Spirodiclofen (0.01) | (a) Spiromesifene (0.01) | (a) Spirotetramate (0.01) | (a) Spiroxamine (0.005) | (a) Tébuconazole (0.005) | (a) Tébufénozide (0.005) |
| (a) Tébufenpyrad (0.005) | (a) Teflubenzuron (0.05) | (a) TEPP (0.01) | (a) Terbacile (0.01) | (a) Terbufos (0.005) | (a) Terbufos-sulfon (0.005) |
| (a) Terbufos-sulfoxide (0.005) | (a) Terbuthylazine (0.01) | (a) Terbutryne (0.005) | (a) Tetraconazole (0.01) | (a) Thiabendazole (0.005) | (a) Thiacloprid (0.005) |
| (a) Thiamethoxam (0.005) | (a) Thiazafluron (0.005) | (a) Thifensulfuron méthyle (0.005) | (a) Thiocarbazil (0.01) | (a) Thiodicarbe (0.005) | (a) Thiofanox (0.01) |
| (a) Thiofanox-Sulfone (0.005) | (a) Thiofanox-Sulfoxid (0.005) | (a) Thiométon (0.05) | (a) Thionazin (0.01) | (a) Thiophanate-éthyl (0.005) | (a) Thiophanate-méthyl (0.005) |
| (a) Triadimefone (0.01) | (a) Triadimenole (0.01) | (a) Triamiphos (0.01) | (a) Triasulfuron (0.005) | (a) Triazamate (0.01) | (a) Triazophos (0.005) |
| (a) Tribenuron methyl (0.01) | (a) Trichlorfon (0.05) | (a) Tricyclazole (0.01) | (a) Tridemorph (0.01) | (a) Trietazine (0.01) | (a) Trifloxystrobine (0.01) |
| (a) Trifloxysulfuron (0.01) | (a) Triflumizol (0.005) | (a) Triflumuron (0.01) | (a) Triflusulfuron-methyl (0.005) | (a) Triforine (0.01) | (a) Trimethacarb 3.4.5- (0.01) |
| (a) Triticonazole (0.005) | (a) Uniconazole (0.005) | (a) Vamidothion (0.005) | (a) Vamidothion-sulfone (0.01) | (a) Vamidothion-sulfoxide (0.01) | (a) Zoxamide (0.01) |

**SIGNATURE**

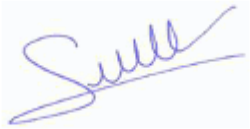

Jérôme Ginet
Business Unit Manager

Rapport validé électroniquement par Jérôme Ginet

**NOTE EXPLICATIVE**

Ce document ne concerne que l'objet soumis à l'essai ; sa reproduction n'est autorisée que sous sa forme intégrale.
Les essais et rapports sont réalisés conformément à nos conditions générales de vente disponibles sur demande.
Pour déclarer ou non la conformité, l'incertitude associée au résultat a été ajoutée ou retranchée de façon à obtenir sans conteste un résultat opposable aux spécifications ou à la réglementation. Elle n'a pas été prise en compte dans le cadre des référentiels qui intègrent déjà les incertitudes de mesures ou sur demande explicite du client.
Les essais sont identifiés par un code de 5 caractères dont la description précise est disponible sur demande.

Les essais identifiés par le code à 2 lettres SF ont été réalisés par le laboratoire SOFIA (Berlin). Le symbole (a) identifie les prestations couvertes par l'accréditation DIN EN ISO/IEC 17025:2005 DAKKS D-PL-19579-02-00.

**Eurofins Analytics France (Nantes)**
Rue Pierre Adolphe Bobierre
BP 42301
F-44323 Nantes Cedex 3
**FRANCE**

Tél. +33 2 51 83 43 40

www.eurofins.fr

SAS au capital de 3 256 700 €
RCS NANTES 423 190 891
SIRET 423 190 891 00022
APE 743 B

Accréditation(s)
N° : 1-2335
N° : 1-2338
Liste
des sites
accrédités
et portée(s)
disponible(s) sur
www.cofrac.fr

SAINT NOLFF - DEPARTEMENT CHIMIE

CS 40234
56011 VANNES CEDEX
FRANCE

Tél : +33 (0)1 71 25 06 06
Mail :

SAFE PF
Route de Saint Bris

89290 AUGY
France

### RAPPORT D'ESSAI FINAL

#### PRODUIT FINI - U8220 V257

| | |
|---|---|
| Date réception client : | 10/08/2017 |
| Date fabrication : | |
| N° lot client : | 17207 |
| Fournisseur : | |
| N° lot fournisseur : | |
| Tonnage : | |
| DLUO : | |

| | |
|---|---|
| Demandeur : | M. BARRAL Renaud |
| N° commande : | |
| N° client : | E8793 |
| N° optim : | |
| N° étude : | |
| Réf. commerciale : | |
| Tiers : | |

Date réception labo : 16/08/2017

Masse brute (g): 2003.33

Observations : CD011344

Commentaires :

*Le code à 2 lettres indique le site Invivo Labs sur lequel a été réalisée l'analyse : CT = site de Chierry, SN = site de Saint-Nolff.*

*L'accréditation du Cofrac atteste de la compétence des laboratoires pour les seuls essais couverts par l'accréditation et qui sont identifiés par le fait qu'ils sont soulignés.*
*Les essais soulignés identifiés CT sont couverts par l'accréditation Cofrac n° 1-2338. Les essais soulignés identifiés SN sont couverts par l'accréditation Cofrac n° 1-2335.*
*(portées disponibles sur www.cofrac.fr)*

*En cas de déclaration de conformité à la spécification, celle-ci ne prend pas en compte l'incertitude associée aux résultats.*
*Si ce rapport fait mention de résultats de mycotoxines, ils sont corrigés du taux de récupération. Ce rapport d'essai ne concerne que l'échantillon soumis à essai.*
*Si ce rapport fait mention de résultats de pesticides, ils ne sont pas corrigés du taux de récupération si celui-ci est compris entre 70 et 120 %*
*« # » : analyse faite plusieurs fois*

*La reproduction de ce rapport n'est autorisée que sous sa forme intégrale.*



*Invivo Labs - Siège social : Talhouët 56250 Saint Nolff - Capital 8 181 400 € - 513 504 399 RCS VANNES - Siret : 513 504 399 00033*

### ANALYSES CHIMIQUES

| Determination | Rés/brut | Rés/sec | Incertitude | Cible | Mini | Maxi | Conforme |
|---|---|---|---|---|---|---|---|
| CENDRES BRUTES (pour dosage minéraux)<br>Méthode interne CEND-H 13/02 adaptée du Règlement CE 152/2009 du 27-01-2009 - SN | 5,2<br>g/100g | | 0,2<br>g/100g | | | | |
| FACTEURS ANTITRYPSIQUES<br>AOCS Ba 12-75 - SN | 1,0<br>UTI/mg | | 0,5<br>UTI/mg | | | | |
| PROTEINES DUMAS (Nx6.25)<br>(#) Méthode interne DUMAS-H 14/02 adaptée de la norme NF EN ISO 16634-1 - décembre 2008 - SN | 18,2<br>g/100g | | 0,5<br>g/100g | | | | |
| CELLULOSE BRUTE<br>(#) Méthode interne CELL-H 14/01 adaptée du Règlement CE 152/2009 du 27-01-2009 - SN | 4,8<br>g/100g | | 0,8<br>g/100g | | | | |
| MATIERES GRASSES BRUTES TOTALES<br>Méthode interne - MGRA-H - Procédé B - SN | 3,2<br>g/100g | | 0,5<br>g/100g | | | | |
| CALCIUM<br>Méthode interne - MINEROL 15 - SN | 7713<br>mg/kg | | 771<br>mg/kg | | | | |
| PHOSPHORE<br>Méthode interne - MINEROL 15 - SN | 6064<br>mg/kg | | 485<br>mg/kg | | | | |
| MINERALISATION MICRO ONDES<br>Méthode interne - ELTRACES-H - CT | Réalisée | | | | | | |
| ARSENIC<br>Méthode interne - ELTRACES-H - CT | 903<br>µg/kg | | 181<br>µg/kg | | | | |
| CADMIUM<br>Méthode interne - ELTRACES-H - CT | 71<br>µg/kg | | 14<br>µg/kg | | | | |
| MERCURE<br>Méthode interne - ELTRACES-H - CT | 16<br>µg/kg | | 6<br>µg/kg | | | | |
| PLOMB<br>Méthode interne - ELTRACES-H - CT | 56<br>µg/kg | | 11<br>µg/kg | | | | |
| NITRITES<br>Méthode interne - NIT - SN | <1,0<br>mg/kg exp en NaNO2 | | | | | | |
| NITRATES<br>Méthode interne - NIT - SN | 15<br>mg/kg exp en NaNO3 | | 3<br>mg/kg exp en NaNO3 | | | | |
| AMIDON ENZYMATIQUE<br>(#) NF V18-121 - mars 1997 abrogée - SN | 38,8<br>g/100g | | 1,9<br>g/100g | | | | |
| SUCRES TOTAUX (exprimés en saccharose)<br>Méthode interne SUCRES-H 14/02 adaptée du Règlement CE 152/2009 du 27-01-2009 - CT | 3,7<br>% de saccharose | | 0,5<br>% de saccharose | | | | |
| B.H.T. (HPLC)<br>Méthode interne - OXAOAC - SN | <5<br>mg/kg | | | | | | |
| B.H.A. (HPLC)<br>Méthode interne - OXAOAC - SN | <5<br>mg/kg | | | | | | |
| GALLATE DE PROPYLE<br>Méthode interne - OXAOAC - SN | <5<br>mg/kg | | | | | | |
| GALLATE D'OCTYLE<br>Méthode interne - OXAOAC - SN | <5<br>mg/kg | | | | | | |
| VITAMINE A<br>Méthode interne VIT_A_E-S - SN | 4,0<br>UI/g | | 1,2<br>UI/g | | | | |
| VITAMINE E (acétate DL-a-tocophérol)<br>Méthode interne VIT_A_E-S - SN | 18,0<br>mg/kg | | 3,6<br>mg/kg | | | | |
| POLYPHENOLS TOTAUX (exp. ac.gallique)<br>ISO 14502-1 - mars 2005 - SN | 2548<br>mg/kg | | 255<br>mg/kg | | | | |
| ACIDE PHYTIQUE<br>Méthode interne - PHYTIC06/00 - SN | 1,0<br>g/100g | | 0,1<br>g/100g | | | | |
| PCDD ET PCDF (TEQ-OMS-avec LQ)<br>Analyse sous traitée | 0,038<br>ng/kg de m. à 12% d'eau | | | | | | |

*Le code à 2 lettres indique le site Invivo Labs sur lequel a été réalisée l'analyse : CT = site de Chierry, SN = site de Saint-Nolff.*

*L'accréditation du Cofrac atteste de la compétence des laboratoires pour les seuls essais couverts par l'accréditation et qui sont identifiés par le fait qu'ils sont soulignés. Les essais soulignés identifiés CT sont couverts par l'accréditation Cofrac n° 1-2338. Les essais soulignés identifiés SN sont couverts par l'accréditation Cofrac n° 1-2335. (portées disponibles sur www.cofrac.fr)*

*En cas de déclaration de conformité à la spécification, celle-ci ne prend pas en compte l'incertitude associée aux résultats.*
*Si ce rapport fait mention de résultats de mycotoxines, ils sont corrigés du taux de récupération. Ce rapport d'essai ne concerne que l'échantillon soumis à essai.*
*Si ce rapport fait mention de résultats de pesticides, ils ne sont pas corrigés du taux de récupération si celui-ci est compris entre 70 et 120 %*
*« # » : analyse faite plusieurs fois*

*La reproduction de ce rapport n'est autorisée que sous sa forme intégrale.*

Accréditation(s)
N° : 1-2335
N° : 1-2338
Liste
des sites
accrédités
et portée(s)
disponible(s) sur
www.cofrac.fr

| Determination | Rés/brut | Rés/sec | Incertitude | Cible | Mini | Maxi | Conforme |
|---|---|---|---|---|---|---|---|
| PCB DL (TEQ-OMS-avec LQ)<br>Analyse sous traitée | 0,067<br>ng/kg de m. à 12% d'eau | | | | | | |
| PCDD/F ET PCB DL (TEQ-OMS-avec LQ)<br>Analyse sous traitée | 0,100<br>ng/kg de m. à 12% d'eau | | | | | | |
| PCB NDL (6 PCBs hors CB118)<br>Analyse sous traitée | 0,270<br>µg/kg de pdt. à 12% d'eau | | | | | | |
| EXTRACTION MATIERE GRASSE POUR ACIDES GRAS<br>Méthode interne EXTRACPG 99 - SN | FAIT | | | | | | |

*Le code à 2 lettres indique le site Invivo Labs sur lequel a été réalisée l'analyse : CT = site de Chierry, SN = site de Saint-Nolff.*

*L'accréditation du Cofrac atteste de la compétence des laboratoires pour les seuls essais couverts par l'accréditation et qui sont identifiés par le fait qu'ils sont soulignés.*
*Les essais soulignés identifiés CT sont couverts par l'accréditation Cofrac n° 1-2338. Les essais soulignés identifiés SN sont couverts par l'accréditation Cofrac n° 1-2335.*
*(portées disponibles sur www.cofrac.fr)*

*En cas de déclaration de conformité à la spécification, celle-ci ne prend pas en compte l'incertitude associée aux résultats.*
*Si ce rapport fait mention de résultats de mycotoxines, ils sont corrigés du taux de récupération. Ce rapport d'essai ne concerne que l'échantillon soumis à essai.*
*Si ce rapport fait mention de résultats de pesticides, ils ne sont pas corrigés du taux de récupération si celui-ci est compris entre 70 et 120 %*
*« # » : analyse faite plusieurs fois*

*La reproduction de ce rapport n'est autorisée que sous sa forme intégrale.*

cofrac ESSAIS ACCREDITATION
Accréditation(s)
N° : 1-2335
N° : 1-2338
Liste
des sites
accrédités
et portée(s)
disponible(s) sur
www.cofrac.fr

### PROFILS ANALYTIQUES

#### ACIDES AMINES TOTAUX (sans tyrosine)

Méthode : Règlement CE 152/2009 du 27-01-2009 - SN

| Determination | Unité | Résultat | Rés. / sec | Incertitude | TX recouvrement | Cible | Maxi | Conforme |
|---|---|---|---|---|---|---|---|---|
| CYSTINE | g/100g | 0,29 | | 0,03 | | | | |
| A.ASPARTIQUE | g/100g | 1,46 | | 0,12 | | | | |
| PROLINE | g/100g | 1,31 | | 0,10 | | | | |
| METHIONINE | g/100g | 0,33 | | 0,03 | | | | |
| THREONINE | g/100g | 0,65 | | 0,05 | | | | |
| SERINE | g/100g | 0,82 | | 0,07 | | | | |
| A.GLUTAMIQUE | g/100g | 3,56 | | 0,28 | | | | |
| GLYCINE | g/100g | 0,95 | | 0,08 | | | | |
| ALANINE | g/100g | 0,86 | | 0,07 | | | | |
| VALINE | g/100g | 0,82 | | 0,07 | | | | |
| ISOLEUCINE | g/100g | 0,69 | | 0,06 | | | | |
| LEUCINE | g/100g | 1,29 | | 0,10 | | | | |
| PHENYLALANINE | g/100g | 0,81 | | 0,07 | | | | |
| LYSINE | g/100g | 0,89 | | 0,07 | | | | |
| HISTIDINE | g/100g | 0,41 | | 0,03 | | | | |
| ARGININE | g/100g | 1,04 | | 0,08 | | | | |
| Total des A.aminés quantifiés | g/100g | 16,17 | | | | | | |

#### SPECTRE DES SUCRES

Méthode : Méthode interne - SUCRES 14 - CT

| Determination | Unité | Résultat | Rés. / sec | Incertitude | TX recouvrement | Cible | Maxi | Conforme |
|---|---|---|---|---|---|---|---|---|
| FRUCTOSE | g/100g | <0,2 | | | | | | |
| GLUCOSE | g/100g | <0,2 | | | | | | |
| SACCHAROSE | g/100g | 1,1 | | 0,2 | | | | |
| LACTOSE | g/100g | <0,4 | | | | | | |
| MALTOSE | g/100g | <0,4 | | | | | | |
| SOMME DES SUCRES | g/100g | 1,1 | | | | | | |

*Le code à 2 lettres indique le site Invivo Labs sur lequel a été réalisée l'analyse : CT = site de Chierry, SN = site de Saint-Nolff.*

*L'accréditation du Cofrac atteste de la compétence des laboratoires pour les seuls essais couverts par l'accréditation et qui sont identifiés par le fait qu'ils sont soulignés.*
*Les essais soulignés identifiés CT sont couverts par l'accréditation Cofrac n° 1-2338. Les essais soulignés identifiés SN sont couverts par l'accréditation Cofrac n° 1-2335.*
*(portées disponibles sur www.cofrac.fr)*

*En cas de déclaration de conformité à la spécification, celle-ci ne prend pas en compte l'incertitude associée aux résultats.*
*Si ce rapport fait mention de résultats de mycotoxines, ils sont corrigés du taux de récupération. Ce rapport d'essai ne concerne que l'échantillon soumis à essai.*
*Si ce rapport fait mention de résultats de pesticides, ils ne sont pas corrigés du taux de récupération si celui-ci est compris entre 70 et 120 %*
*« # » : analyse faite plusieurs fois*

*La reproduction de ce rapport n'est autorisée que sous sa forme intégrale.*

#### ISOFLAVONES DE SOJA

Méthode : Méthode interne - SOJISOFLAV 06/00 - SN

| Determination | Unité | Résultat | Rés. / sec | Incertitude | TX recouvrement | Cible | Maxi | Conforme |
|---|---|---|---|---|---|---|---|---|
| DAIDZIN | mg/kg | 55,9 | | | | | | |
| GLYCITIN | mg/kg | <0,5 | | | | | | |
| GENISTIN | mg/kg | 68,1 | | | | | | |
| DAIDZÉIN | mg/kg | 6,0 | | | | | | |
| GLYCITÉIN | mg/kg | 6,5 | | | | | | |
| GENISTÉIN | mg/kg | 4,3 | | | | | | |
| ISOFLAVONES totaux | mg/kg | 140,8 | | | | | | |

#### MYCOTOXINES

Méthode : Méthode interne - MULTIMYC3 Protocole A - CT

| Determination | Unité | Résultat | Rés. / sec | Incertitude | TX recouvrement | Cible | Maxi | Conforme |
|---|---|---|---|---|---|---|---|---|
| AFLATOXIN B1 | µg/kg | <0,5 | | | 103 | | | |
| AFLATOXIN B2 | µg/kg | <0,5 | | | 91 | | | |
| AFLATOXINE G1 | µg/kg | <0,5 | | | 86 | | | |
| AFLATOXINE G2 | µg/kg | <0,5 | | | 69 | | | |
| SOMME DES AFLATOXINES (B1+B2+G1+G2) | µg/kg | <2,0 | | | | | | |
| OCHRATOXINE A | µg/kg | 2,45 | | 1,23 | 64 | | | |
| DEOXYNIVALENOL | µg/kg | 339 | | 68 | 85 | | | |
| SOMME (15A DON & 3A DON) | µg/kg | <50 | | | 91 | | | |
| NIVALENOL | µg/kg | <50 | | | 67 | | | |
| FUSARENONE X | µg/kg | <50 | | | 88 | | | |
| ZEARALENONE | µg/kg | 25 | | 6 | 79 | | | |
| T2 TOXINE | µg/kg | <50 | | | 109 | | | |
| HT2 TOXINE | µg/kg | <50 | | | 89 | | | |
| SOMME T2 + HT2 | µg/kg | <100 | | | | | | |
| DIACETOXYSCIRPENOL | µg/kg | <50 | | | 77 | | | |
| NEOSOLANIOL | µg/kg | <50 | | | 77 | | | |
| FUMONISINE B1 | µg/kg | <10 | | | 88 | | | |
| FUMONISINE B2 | µg/kg | <10 | | | 66 | | | |

*Le code à 2 lettres indique le site Invivo Labs sur lequel a été réalisée l'analyse : CT = site de Chierry, SN = site de Saint-Nolff.*

*L'accréditation du Cofrac atteste de la compétence des laboratoires pour les seuls essais couverts par l'accréditation et qui sont identifiés par le fait qu'ils sont soulignés. Les essais soulignés identifiés CT sont couverts par l'accréditation Cofrac n° 1-2338. Les essais soulignés identifiés SN sont couverts par l'accréditation Cofrac n° 1-2335. (portées disponibles sur www.cofrac.fr)*

*En cas de déclaration de conformité à la spécification, celle-ci ne prend pas en compte l'incertitude associée aux résultats.*
*Si ce rapport fait mention de résultats de mycotoxines, ils sont corrigés du taux de récupération. Ce rapport d'essai ne concerne que l'échantillon soumis à essai.*
*Si ce rapport fait mention de résultats de pesticides, ils ne sont pas corrigés du taux de récupération si celui-ci est compris entre 70 et 120 %*
*« # » : analyse faite plusieurs fois*

*La reproduction de ce rapport n'est autorisée que sous sa forme intégrale.*

Accréditation(s)
N° : 1-2335
N° : 1-2338
Liste
des sites
accrédités
et portée(s)
disponible(s) sur
www.cofrac.fr

### MYCOTOXINES

#### Méthode : Méthode interne - MULTIMYC3 Protocole A - CT

| Determination | Unité | Résultat | Rés. / sec | Incertitude | TX recouvrement | Cible | Maxi | Conforme |
|---|---|---|---|---|---|---|---|---|
| FUMONISINE B3 | µg/kg | <10 | | | 75 | | | |
| SOMME DES FUMONISINES (B1+B2) | µg/kg | <20 | | | | | | |

*Le code à 2 lettres indique le site Invivo Labs sur lequel a été réalisée l'analyse : CT = site de Chierry, SN = site de Saint-Nolff.*

*L'accréditation du Cofrac atteste de la compétence des laboratoires pour les seuls essais couverts par l'accréditation et qui sont identifiés par le fait qu'ils sont soulignés.*
*Les essais soulignés identifiés CT sont couverts par l'accréditation Cofrac n° 1-2338. Les essais soulignés identifiés SN sont couverts par l'accréditation Cofrac n° 1-2335.*
*(portées disponibles sur www.cofrac.fr)*

*En cas de déclaration de conformité à la spécification, celle-ci ne prend pas en compte l'incertitude associée aux résultats.*
*Si ce rapport fait mention de résultats de mycotoxines, ils sont corrigés du taux de récupération. Ce rapport d'essai ne concerne que l'échantillon soumis à essai.*
*Si ce rapport fait mention de résultats de pesticides, ils ne sont pas corrigés du taux de récupération si celui-ci est compris entre 70 et 120 %*
*« # » : analyse faite plusieurs fois*
*La reproduction de ce rapport n'est autorisée que sous sa forme intégrale.*

*Invivo Labs - Siège social : Talhouët 56250 Saint Nolff - Capital 8 181 400 € - 513 504 399 RCS VANNES - Siret : 513 504 399 00033*

Accréditation(s)
N° : 1-2335
N° : 1-2338
Liste des sites accrédités et portée(s) disponible(s) sur www.cofrac.fr

### PROFIL D'ESTERS D'ACIDES GRAS BRUT AVEC TRANS

NF EN ISO 12966-2 - Juin 2011 / NF EN ISO 12966-4 - Août 2015

#### BILAN

| | Composition en | % relatif | mg/100g | val. usuelle | Mini | Maxi |
|---|---|---|---|---|---|---|
| **Acides Gras Saturés** | | | | | | |
| Total acides gras saturés | AGS | 20,2 | 511 | | | |
| Acide palmitique | C16:0 | 16,5 | 418 | | | |
| Acides stéarique | C18:0 | 2,0 | 51 | | | |
| **Acides Gras Insaturés** | | | | | | |
| Total acides gras insaturés | AGI | 79,8 | 1967 | | | |
| Acide oléïque et isomères | C18:1 | 21,9 | 544 | | | |
| Total acides gras mono-insaturés | AGMI | 24,9 | 618 | | | |
| Total acides gras poly-insaturés | AGPI | 54,9 | 1349 | | | |
| **Oméga 3** | | | | | | |
| Total acides gras omega 3 | n-3 | 6,4 | 155 | | | |
| Acide alpha-linolénique (ALA) | ALA | 4,2 | 103 | | | |
| Acide eicosapentaènoïque (EPA) | EPA | 0,6 | 15 | | | |
| Acide docosahexaènoïque (DHA) | DHA | 1,0 | 24 | | | |
| Acide docosapentaènoïque (DPA) | DPA | 0,3 | <10 | | | |
| **Oméga 6** | | | | | | |
| Total acides gras omega 6 | n-6 | 47,8 | 1177 | | | |
| Acide linoléique (LA) | LA | 47,5 | 1170 | | | |
| **Acides Gras Trans** | | | | | | |
| Total acides gras trans (hors CLA) | AGtr tx | 1,1 | 28 | | | |
| Total 18:1 trans | | 0,5 | 12 | | | |
| **CLA** | | | | | | |
| CLA totaux | CLA | nd | nd | | | |

#### RAPPORTS ET CRITERES SPECIFIQUES

| | | | val. usuelle | Mini | Maxi |
|---|---|---|---|---|---|
| LA / ALA | LA/ALA | 11,3 | | | |
| n-6 / n-3 | n-6/n-3 | 7,5 | | | |
| C18:1 / C16:0 | C18:1/C16:0 | 1,3 | | | |
| AGS / n-3 | AGS/n-3 | 3,2 | | | |
| C16:0 / AGS | C16:0/AGS | 0,8 | | | |
| C16:0 / ALA | C16:0/ALA | 3,9 | | | |
| AGPI / ALA | AGPI/ALA | 13,1 | | | |
| C18:1 tr11 / C18:1 tr10 | C18:1tr11/C18:1tr10 | 1,0 | | | |
| AGS/ALA | AGS/ALA | 4,8 | | | |

| | Composition en | % relatif | mg/100g | val. usuelle | Mini | Maxi |
|---|---|---|---|---|---|---|
| AGMI+C18:0-C16:0 | | 10,4 | 251 | | | |
| Acides Gras sur produit sec | (g d'AG/kg de MS) | | | | | |
| ALA sur produit sec | (g d'ALA/kg de MS) | | | | | |

*Le code à 2 lettres indique le site Invivo Labs sur lequel a été réalisée l'analyse : CT = site de Chierry, SN = site de Saint-Nolff.*

*L'accréditation du Cofrac atteste de la compétence des laboratoires pour les seuls essais couverts par l'accréditation et qui sont identifiés par le fait qu'ils sont soulignés. Les essais soulignés identifiés CT sont couverts par l'accréditation Cofrac n° 1-2338. Les essais soulignés identifiés SN sont couverts par l'accréditation Cofrac n° 1-2335. (portées disponibles sur www.cofrac.fr)*

*En cas de déclaration de conformité à la spécification, celle-ci ne prend pas en compte l'incertitude associée aux résultats.*
*Si ce rapport fait mention de résultats de mycotoxines, ils sont corrigés du taux de récupération. Ce rapport d'essai ne concerne que l'échantillon soumis à essai.*
*Si ce rapport fait mention de résultats de pesticides, ils ne sont pas corrigés du taux de récupération si celui-ci est compris entre 70 et 120 %*
*« # » : analyse faite plusieurs fois*

*La reproduction de ce rapport n'est autorisée que sous sa forme intégrale.*

Accréditation(s)
N° : 1-2335
N° : 1-2338
Liste
des sites
accrédités
et portée(s)
disponible(s) sur
www.cofrac.fr

| Composition en | | % relatif | mg/100g | val. usuelle | Mini | Maxi |
|---|---|---|---|---|---|---|
| Acide butyrique | C 4:0 | nd | nd | | | |
| Acide valérique | C 5:0 | nd | nd | | | |
| Acide caproïque | C 6:0 | nd | nd | | | |
| Acide heptanoïque | C 7:0 | nd | nd | | | |
| Acide caprylique | C 8:0 | nd | nd | | | |
| Acide nonaïque | C 9:0 | nd | nd | | | |
| Acide caprique | C 10:0 | nd | nd | | | |
| Acide caproléique | C 10:1 | nd | nd | | | |
| Acide undécanoïque | C 11:0 | nd | nd | | | |
| Acide undécénoïque | C 11:1 | nd | nd | | | |
| Acide laurique | C 12:0 | 0,1 | <10 | | | |
| Acide lauroléique | C 12:1 | nd | nd | | | |
| Acide 11-méthyl dodécanoïque | C 13:0 iso | nd | nd | | | |
| Acide 10-méthyl dodécanoïque | C 13:0 anteiso | nd | nd | | | |
| | *total_C13:0* | nd | nd | | | |
| Acide isomyristique | C 14:0 iso | nd | nd | | | |
| Acide myristique | C 14:0 | 0,6 | 15 | | | |
| | *total_C14:0* | 0,6 | 15 | | | |
| Acide myristoléique | C 14:1 | nd | nd | | | |
| Acide 13-méthyl tétradécanoïque | C 15:0 iso | nd | nd | | | |
| Acide 12-méthyl tétradécanoïque | C 15:0 anteiso | nd | nd | | | |
| Acide pentadécanoïque | C 15:0 | 0,1 | <10 | | | |
| | *total_C15:0* | 0,1 | <10 | | | |
| Acide cis-10-pentadécènoïque | C 15:1 n-5 | nd | nd | | | |
| Acide pentadécénoïque | C 15:1 | nd | nd | | | |
| | *total_C15:1* | nd | nd | | | |
| Acide isopalmitique | C 16:0 iso | nd | nd | | | |
| Acide palmitique | C 16:0 | 16,5 | 418 | | | |
| | *total_C16:0* | 16,5 | 418 | | | |
| Acide hypogéïque | C 16:1 n-9 | 0,1 | <10 | | | |
| Acide palmitoléïque | C 16:1 n-7 | 0,6 | 16 | | | |
| Acide hexadécènoïque (autres isomères) | C 16:1 | <0,05 | <10 | | | |
| | *total_C16:1* | 0,7 | 19 | | | |
| Acide hexadécadiènoïque | C 16:2 | <0,05 | <10 | | | |
| Acide hexadécatriènoïque | C 16:3 | nd | nd | | | |
| Acide hexadécatétraènoïque | C 16:4 | nd | nd | | | |


*La reproduction de ce rapport n'est autorisée que sous sa forme intégrale.*

Accréditation(s)
N° : 1-2335
N° : 1-2338
Liste des sites accrédités et portée(s) disponible(s) sur www.cofrac.fr

| Composition en | | %relatif | mg/100g | val. usuelle | Mini | Maxi |
|---|---|---|---|---|---|---|
| Acide isomargarique | C17:0 iso | nd | nd | | | |
| Acide 14-méthyl hexadécanoïque | C17:0 anteiso | nd | nd | | | |
| Acide margarique | C17:0 | 0,1 | <10 | | | |
| | *total_C17:0* | 0,1 | <10 | | | |
| Acide 14-méthyl 8-hexadécènoïque | C17:1 anteiso | <0,05 | <10 | | | |
| Acide heptadécènoïque | C17:1 | 0,1 | <10 | | | |
| | *total_C17:1* | 0,1 | <10 | | | |
| Acide isostéarique | C18:0 iso | nd | nd | | | |
| Acide stéarique | C18:0 | 2,0 | 51 | | | |
| | *total_C18:0* | 2,0 | 51 | | | |
| Acide trans-4-octadécènoïque | C18:1 tr4 | nd | nd | | | |
| Acide trans-5-octadécènoïque | C18:1 tr5 | nd | nd | | | |
| Acide trans-(6-8)-octadécènoïque | C18:1 tr6-8 | nd | nd | | | |
| Acide élaidique | C18:1 tr9 | 0,3 | <10 | | | |
| Acide trans-10-octadécènoïque | C18:1 tr10 | 0,1 | <10 | | | |
| Acide trans-vaccénique | C18:1 tr11 | 0,1 | <10 | | | |
| Acide trans-12-octadécènoïque | C18:1 tr12 | nd | nd | | | |
| | *total_C18:1tr* | 0,5 | 12 | | | |
| Acide oléique * | C18:1 c9 | 20,0 | 496 | | | |
| Acide cis-10-octadécènoïque * | C18:1 c10 | nd | nd | | | |
| Acide cis-vaccénique | C18:1 c11 | 1,3 | 33 | | | |
| Acide cis-12-octadécènoïque | C18:1 c12 | 0,1 | <10 | | | |
| Acide cis-13-octadécènoïque | C18:1 c13 | <0,05 | <10 | | | |
| Acide cis-14-octadécènoïque | C18:1 c14 | nd | nd | | | |
| Acide cis-15-octadécènoïque | C18:1 c15 | nd | nd | | | |
| Acide cis-16-octadécènoïque | C18:1 c16 | nd | nd | | | |
| | *total_C18:1cis* | 21,4 | 532 | | | |
| | *total_C18:1* | 21,9 | 544 | | | |
| Acide linolélaidique | C18:2 n-6 tr | 0,1 | <10 | | | |
| Acide octadécadiènoïque (isomère cis-trans) | C18:2 ct | 0,3 | <10 | | | |
| Acide octadécadiènoïque (isomère trans-cis) | C18:2 tc | 0,2 | <10 | | | |
| | *total_C18:2tr* | 0,6 | 15 | | | |
| Acide linoléique (LA) | C18:2 n-6 | 47,5 | 1170 | | | |
| Acide octadécadiènoïque (autres isomères cis) | C18:2 cis | nd | nd | | | |
| | *total_C18:2cis* | 47,5 | 1170 | | | |

*Le code à 2 lettres indique le site Invivo Labs sur lequel a été réalisée l'analyse : CT = site de Chierry, SN = site de Saint-Nolff.*

*L'accréditation du Cofrac atteste de la compétence des laboratoires pour les seuls essais couverts par l'accréditation et qui sont identifiés par le fait qu'ils sont soulignés.*
*Les essais soulignés identifiés CT sont couverts par l'accréditation Cofrac n° 1-2338. Les essais soulignés identifiés SN sont couverts par l'accréditation Cofrac n° 1-2335.*
*(portées disponibles sur www.cofrac.fr)*

*En cas de déclaration de conformité à la spécification, celle-ci ne prend pas en compte l'incertitude associée aux résultats.*
*Si ce rapport fait mention de résultats de mycotoxines, ils sont corrigés du taux de récupération. Ce rapport d'essai ne concerne que l'échantillon soumis à essai.*
*Si ce rapport fait mention de résultats de pesticides, ils ne sont pas corrigés du taux de récupération si celui-ci est compris entre 70 et 120 %*
*« # » : analyse faite plusieurs fois*

*La reproduction de ce rapport n'est autorisée que sous sa forme intégrale.*

cofrac ESSAIS ACCREDITATION
Accréditation(s)
N° : 1-2335
N° : 1-2338
Liste des sites accrédités et portée(s) disponible(s) sur www.cofrac.fr

| Composition en | | % relatif | mg/100g | val. usuelle | Mini | Maxi |
|---|---|---|---|---|---|---|
| Acide ruménique (CLA) | CLA c9tr11 | nd | nd | | | |
| Acide linoléique conjugué (CLA) | CLA tr10c12 | nd | nd | | | |
| Acide linoléique conjugué (CLA, isomères) | CLA | nd | nd | | | |
| | *total_CLA* | nd | nd | | | |
| | *total_C18:2* | 48,1 | 1185 | | | |
| Acide octadécatriènoïque (isomères trans) | C18:3 tr | 0,1 | <10 | | | |
| Acide octadécatriènoïque (isomères cis) | C18:3 cis | nd | nd | | | |
| Acide gamma-linolénique (GLA) | C18:3 n-6 | nd | nd | | | |
| Acide alpha-linolénique (ALA) | C18:3 n-3 | 4,2 | 103 | | | |
| | *total_C18:3* | 4,3 | 105 | | | |
| Acide stéaridonique | C18:4 n-3 | 0,1 | <10 | | | |
| Acide octadécatétraènoïque (autres isomères) | C18:4 | nd | nd | | | |
| | *total_C18:4* | 0,1 | <10 | | | |
| Acide nonadécanoïque | C19:0 | nd | nd | | | |
| Acide nonadécènoïque | C19:1 | nd | nd | | | |
| Acide arachidique | C20:0 | 0,3 | <10 | | | |
| Acide cis-5-eicosènoïque | C20:1 n-15 | nd | nd | | | |
| Acide cis-8-eicosènoïque | C20:1 n-12 | nd | nd | | | |
| Acide gadoleïque | C20:1 n-9 | 1,3 | 31 | | | |
| Acides gadoleïque et isomères | C20:1 | nd | nd | | | |
| | *total_C20:1* | 1,3 | 31 | | | |
| Acide eicosadiènoïque | C20:2 n-6 | 0,2 | <10 | | | |
| Acide eicosadiènoïque (autres isomères) | C20:2 | nd | nd | | | |
| | *total_C20:2* | 0,2 | <10 | | | |
| Acide de Mead | C20:3 n-9 | nd | nd | | | |
| Acide eicosatriènoïque (DGLA) | C20:3 n-6 | nd | nd | | | |
| Acide eicosatriènoïque (DALA) | C20:3 n-3 | 0,1 | <10 | | | |
| | *total_C20:3* | 0,1 | <10 | | | |
| Acide arachidonique (AA) | C20:4 n-6 | 0,1 | <10 | | | |
| Acide eicosatétraènoïque | C20:4 n-3 | 0,1 | <10 | | | |
| | *total_C20:4* | 0,2 | <10 | | | |
| Acide eicosapentaènoïque (EPA) | C20:5 n-3 | 0,6 | 15 | | | |
| Acide hénéiconanoïque | C21:0 | nd | nd | | | |
| Acide béhénique | C22:0 | 0,2 | <10 | | | |

*Le code à 2 lettres indique le site Invivo Labs sur lequel a été réalisée l'analyse : CT = site de Chierry, SN = site de Saint-Nolff.*

*L'accréditation du Cofrac atteste de la compétence des laboratoires pour les seuls essais couverts par l'accréditation et qui sont identifiés par le fait qu'ils sont soulignés.*
*Les essais soulignés identifiés CT sont couverts par l'accréditation Cofrac n° 1-2338. Les essais soulignés identifiés SN sont couverts par l'accréditation Cofrac n° 1-2335.*
*(portées disponibles sur www.cofrac.fr)*

*En cas de déclaration de conformité à la spécification, celle-ci ne prend pas en compte l'incertitude associée aux résultats.*
*Si ce rapport fait mention de résultats de mycotoxines, ils sont corrigés du taux de récupération. Ce rapport d'essai ne concerne que l'échantillon soumis à essai.*
*Si ce rapport fait mention de résultats de pesticides, ils ne sont pas corrigés du taux de récupération si celui-ci est compris entre 70 et 120 %*
*« # » : analyse faite plusieurs fois*
*La reproduction de ce rapport n'est autorisée que sous sa forme intégrale.*
*Invivo Labs - Siège social : Talhouët 56250 Saint Nolff - Capital 8 181 400 € - 513 504 399 RCS VANNES - Siret : 513 504 399 00033*

cofrac ESSAIS ACCREDITATION
Accréditation(s) N° : 1-2335 N° : 1-2338 Liste des sites accrédités et portée(s) disponible(s) sur www.cofrac.fr

| Composition en | | % relatif | mg/100g | val. usuelle | Mini | Maxi |
|---|---|---|---|---|---|---|
| Acide cétoléique | C22:1 n-11 | 0,6 | 15 | | | |
| Acide érucique | C22:1 n-9 | 0,1 | <10 | | | |
| Acide docosènoïque | C22:1 n-7 | nd | nd | | | |
| | *total_C22:1* | 0,7 | 18 | | | |
| Acide docosadiènoïque (n-6) | C22:2 n-6 | nd | nd | | | |
| Acide docosadiènoïque | C22:2 | nd | nd | | | |
| | *total_C22:2* | nd | nd | | | |
| Acide docosatriènoïque (n-6) | C22:3 n-6 | nd | nd | | | |
| Acide docosatriènoïque (n-3) | C22:3 n-3 | nd | nd | | | |
| | *total_C22:3* | nd | nd | | | |
| Acide docosatétraènoïque (n-6) | C22:4 n-6 | nd | nd | | | |
| Acide docosatétraènoïque (n-3) | C22:4 n-3 | nd | nd | | | |
| | *total_C22:4* | nd | nd | | | |
| Acide docosapentaènoïque (n-6) | C22:5 n-6 | nd | nd | | | |
| Acide docosapentaènoïque (DPA) | C22:5 n-3 | 0,3 | <10 | | | |
| | *total_C22:5* | 0,3 | <10 | | | |
| Acide docosahexaènoïque (DHA) | C22:6 n-3 | 1,0 | 24 | | | |
| Acide tricosanoïque | C23:0 | 0,1 | <10 | | | |
| Acide lignocérique | C24:0 | 0,2 | <10 | | | |
| Acide nervonique | C24:1 n-9 | 0,2 | <10 | | | |

nd : pic non présent sur le chromatogramme
NR : non recherché par cette méthode
* : Coélution des isomères
La somme des esters méthyliques d'acides gras correspond à la somme des esters méthyliques d'acides gras identifiés.
La quantification des esters méthyliques d'acides gras est déterminée par étalonnage interne.
Un facteur de correction est utilisé pour le calcul des esters méthyliques d'acides gras de C4 à C10.
Les esters méthyliques d'acides gras de C4 à C7 ne sont pas dans le domaine d'application de la norme. Ils sont toutefois inclus dans la somme des Acides Gras Saturés (AGS).

Incertitude en relatif : 8 % de la teneur relative avec une valeur mini de 0,5 et maxi de 3,5.
Incertitude en absolu : 10 mg/100g pour les teneurs < 50mg/100g, pour les teneurs >= 50mg/100g : 12 % de la valeur.

Conclusion :

Validé le : 04-10-17

M. KERRAND Jérémie

Superviseur

Accréditation N°1-1531 PORTEE DISPONIBLE SUR WWW.COFRAC.FR

| RAPPORT D'ESSAI | ANALYSE DES PCDD ET PCDF, DES PCB "type dioxine" ET DES PCB indicateurs |
|---|---|

L'essai LSE17-122614-1 a été réalisé à la demande de Date : 07/09/2017

INVIVO LABS
Contact
TALHOUET ST NOLFF
CS 40234
VANNES 56011

Code essai CARSO-LSEH : LSE17-122614-1
Référence client dossier : Réf envoi 170817-027

#### OBJET DE L'ESSAI

L'objet de ce rapport d'essai référencé sous le code d'essai LSE17-122614 est l'analyse des PCDD et PCDF, et des PCB "type dioxine" et indicateurs.

#### INFORMATIONS SPECIFIQUES A L'ESSAI

| Description | Information |
|---|---|
| Date de réception des échantillons | LSE1708-48971 18/08/2017 |
| Méthode(s) d'analyse - PCDD/F | LSE1708-48971 MET009 |
| Méthode(s) d'analyse - PCB | MET038 |
| Instrument de mesure HRGC/HRMS<br>Volume injecté en micro-litres | Autospec ULTIMA (Waters)<br>1 à 3 microlitres |
| Volume final | 25-50 microlitres |
| Observations spécifiques à l'essai : | LSE1708-48971 Rien à signaler |

Les méthodes internes sont la traduction technique des référentiels suivants : normes EPA 1613, EPA 1668 et EN 16215 (échantillons agro-alimentaires).
Dans le cas des échantillons agro-alimentaires, les méthodes d'analyse sont conformes aux critères énoncés dans le règlement (UE) n° 771/2017 de la commission du 3 mai 2017 (alimentation animale) et dans le règlement (UE) n° 644/2017 de la commission du 5 avril 2017 (alimentation humaine).

Les prélèvements ont été réalisés par le client.

#### RESULTATS

Les résultats résumés dans les tableaux ci-dessous sont obtenus en considérant les valeurs des différents congénères au-dessous de la limite de quantification comme étant égales à la limite de quantification (résultat upperbound).

Les résultats complets sont rapportés dans la deuxième partie du rapport.

**Résumé des résultats en PCDD/F-TEQ**

| Référence client échantillon | Référence CARSO-LSEH | PCDD/F-TEQ | Unité | | Incertitude élargie (k=2) +/-15% |
|---|---|---|---|---|---|
| 17SA052492 - PRODUIT FINI - U8220 V257 | **LSE1708-48971** | 0.038 | ng/kg | de matière à 12% eau (TEF OMS 2005) | 0.006 |

**Résumé des résultats en PCB-TEQ (PCB "Dioxin-like")**

| Référence client échantillon | Référence CARSO-LSEH | PCB-TEQ | Unité | | Incertitude élargie (k=2) +/-15% |
|---|---|---|---|---|---|
| 17SA052492 - PRODUIT FINI - U8220 V257 | **LSE1708-48971** | 0.067 | ng/kg | de matière à 12% eau (TEF OMS 2005) | 0.010 |


Société par action simplifiée au capital de 2 283 622,30 € - RCS Lyon B 410 545 313 - SIRET 410 545 313 00042 - APE 7120B — N° TVA: FR 82 410 545 313
Siège social et laboratoire : 4, avenue Jean Moulin — CS 30228 - 69633 VENISSIEUX CEDEX - Tél : (33) 04 72 76 16 16 - Fax : (33) 04 78 72 35 03
Site web : www.groupecarso.com -

**Résumé des résultats en PCDD/F-PCB-TEQ (PCDD/F + PCB "Dioxin-like")**

| Référence client échantillon | Référence CARSO-LSEH | PCDD/F-PCB-TEQ | Unité | | Incertitude élargie (k=2) +/-15% |
|---|---|---|---|---|---|
| 17SA052492 - PRODUIT FINI - U8220 V257 | **LSE1708-48971** | 0.10 | ng/kg | de matière à 12% eau (TEF OMS 2005) | 0.02 |

**Résumé des résultats en PCB (6 PCBs  hors PCB118)**

| Référence client échantillon | Référence CARSO-LSEH | PCB NDL (6 PCBs hors PCB118) | Unité | | Incertitude élargie (k=2) +/-15% |
|---|---|---|---|---|---|
| 17SA052492 - PRODUIT FINI - U8220 V257 | **LSE1708-48971** | 0.27 | µg/kg | de matière à 12% eau | 0.04 |

**Conformités aux réglementations**

| Référence client échantillon | Référence CARSO-LSEH | Conformité |
|---|---|---|
| 17SA052492 - PRODUIT FINI - U8220 V257 | **LSE1708-48971** | Conforme au règlement (UE) N° 277/2012 du 28 Mars 2012. |

Pour déclarer ou non la conformité à la spécification, l'incertitude associée au résultat a été prise en compte.

La déclaration de conformité concerne uniquement les composés demandés par le client pour lesquels les réglementations en vigueur s'appliquent.

Dans le cas d'échantillons contenant de la matière grasse, le pourcentage est déterminé par pesée.

Dans le cas d'échantillons dont la teneur en eau est communiquée, cette dernière est déterminée par dessication puis pesée de la perte de poids de l'échantillon.

La reproduction de ce document n'est autorisée que sous la forme de fac-similé photographique intégral.
Il comporte 4 pages.

Le rapport établi ne concerne que les échantillons soumis à l'essai.
L'accréditation du COFRAC atteste de la compétence des laboratoires pour les seuls essais couverts par l'accréditation, identifiés par le symbole #.

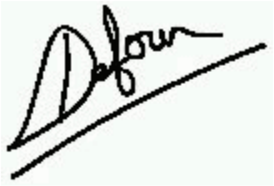

Stéphanie DEFOUR

Responsable de Laboratoire


Société par action simplifiée au capital de 2 283 622,30 € - RCS Lyon B 410 545 313 - SIRET 410 545 313 00042 - APE 7120B — N° TVA: FR 82 [illegible] 313
Siège social et laboratoire : 4, avenue Jean Moulin — CS 30228 - 69633 VENISSIEUX CEDEX - Tél : (33) 04 72 76 16 16 - Fax : (33) 04 78 72 35 03
Site web : www.groupecarso.com -

**Essai LSE17-122614 : Echantillon LSE1708-48971**

Client INVIVO LABS

Date : 07/09/2017

Référence client échantillon 17SA052492 - PRODUIT FINI - U8220 V257

Teneur en eau (%) : 12.71
Teneur en matière grasse (%) : 3.05
Matière à 12% eau (g) : 19.79

Date de début d'analyse : 18/08/2017
Fichiers HRGC/HRMS-PCDD/F : 05SEPV38 - PCB: 05SEPU36 05SEPU36 05SEPV38

| | ng/kg de matière à 12% eau | Taux de récupération % | Cofrac |
|---|---|---|---|
| 2,3,7,8-TeCDD | <0.0051 | 74 | # |
| 1,2,3,7,8-PeCDD | <0.0126 | 86 | # |
| 1,2,3,4,7,8-HxCDD | <0.0126 | 71 | # |
| 1,2,3,6,7,8-HxCDD | <0.0126 | 71 | # |
| 1,2,3,7,8,9-HxCDD | <0.0126 | | # |
| 1,2,3,4,6,7,8-HpCDD | <0.1010 | 81 | # |
| OcCDD | 0.2184 | 73 | # |
| 2,3,7,8-TeCDF | 0.0487 | 60 | # |
| 1,2,3,7,8-PeCDF | <0.0126 | 70 | # |
| 2,3,4,7,8-PeCDF | 0.0153 | 75 | # |
| 1,2,3,4,7,8-HxCDF | <0.0126 | 63 | # |
| 1,2,3,6,7,8-HxCDF | <0.0126 | 66 | # |
| 2,3,4,6,7,8-HxCDF | <0.0126 | 70 | # |
| 1,2,3,7,8,9-HxCDF | <0.0126 | 70 | # |
| 1,2,3,4,6,7,8-HpCDF | <0.0253 | 72 | # |
| 1,2,3,4,7,8,9-HpCDF | <0.0253 | 74 | # |
| OcCDF | <0.0505 | 73 | # |
| PCDD/F-TEQ lower bound (TEF OMS 2005) | 0.0095 | | # |
| PCDD/F-TEQ medium bound (TEF OMS 2005) | 0.024 | | # |
| PCDD/F-TEQ upper bound (TEF OMS 2005) | 0.038 | | # |
| PCB 77 | 2.8970 | 49 | # |
| PCB 81 | <0.1010 | 46 | # |
| PCB 105 | 11.6385 | 102 | # |
| PCB 114 | <2.0202 | 102 | # |
| PCB 118 | 42.1377 | 95 | # |
| PCB 123 | <2.0202 | 96 | # |
| PCB 126 | 0.6144 | 68 | # |
| PCB 156 | 4.6307 | 93 | # |
| PCB 157 | <2.0202 | 93 | # |
| PCB 167 | 3.4084 | 84 | # |
| PCB 169 | <0.1010 | 91 | # |
| PCB 189 | <2.0202 | 44 | # |
| PCB-TEQ lower bound (TEF OMS 2005) | 0.064 | | # |
| PCB-TEQ medium bound (TEF OMS 2005) | 0.065 | | # |
| PCB-TEQ upper bound (TEF OMS 2005) | 0.067 | | # |
| PCDD/F-PCB-TEQ lower bound (TEF OMS 2005) | 0.073 | | # |
| PCDD/F-PCB-TEQ medium bound (TEF OMS 2005) | 0.089 | | # |
| PCDD/F-PCB-TEQ upper bound (TEF OMS 2005) | 0.10 | | # |
| PCB 28 | <10.10 | 69 | # |
| PCB 52 | 14.77 | 66 | # |
| PCB 101 | 29.98 | 67 | # |
| PCB 138 | 67.62 | 94 | # |
| PCB 153 | 117.71 | 74 | # |
| PCB 180 | 31.67 | 78 | # |
| | µg/kg de matière à 12% eau | | |
| PCB NDL (6 PCBs hors PCB118) lower bound | 0.26 | | # |
| PCB NDL (6 PCBs hors PCB118) medium bound | 0.27 | | # |
| PCB NDL (6 PCBs hors PCB118) upper bound | 0.27 | | # |

Lorsque la concentration en analyte est précédée de « < », le résultat communiqué correspond à la limite de quantification (LOQ).

**Légende :** LOQ = Limite de quantification
Lower bound : La valeur 0 est affectée aux congénères <LOQ
Medium bound : La valeur ½ LOQ est affectée aux congénères <LOQ
Upper bound : La valeur de leur LOQ est affectée aux congénères <LOQ

Dans le cas des échantillons agro-alimentaires, la limite de quantification est telle que définie dans l'annexe I du règlement (UE) n° 644/2017. Il s'agit de la concentration de l'analyte dans l'extrait qui produit une réponse instrumentale aux deux ions suivis avec un rapport S/B (signal sur bruit) de 3:1 pour le signal le moins intense et remplit les critères d'identification tels que définis dans la méthode EPA 1613, Révision B.


Société par action simplifiée au capital de 2 283 622,30 € - RCS Lyon B 410 545 313 - SIRET 410 545 313 00042 - APE 7120B — N° TVA: FR 82 410 545 313
Siège social et laboratoire : 4, avenue Jean Moulin — CS 30228 - 69633 VENISSIEUX CEDEX - Tél : (33) 04 72 76 16 16 - Fax : (33) 04 78 72 35 03
Site web : www.groupecarso.com -

Concentration des 17 congénères toxiques

Référence CARSO-LSEHL : LSE1708-48971

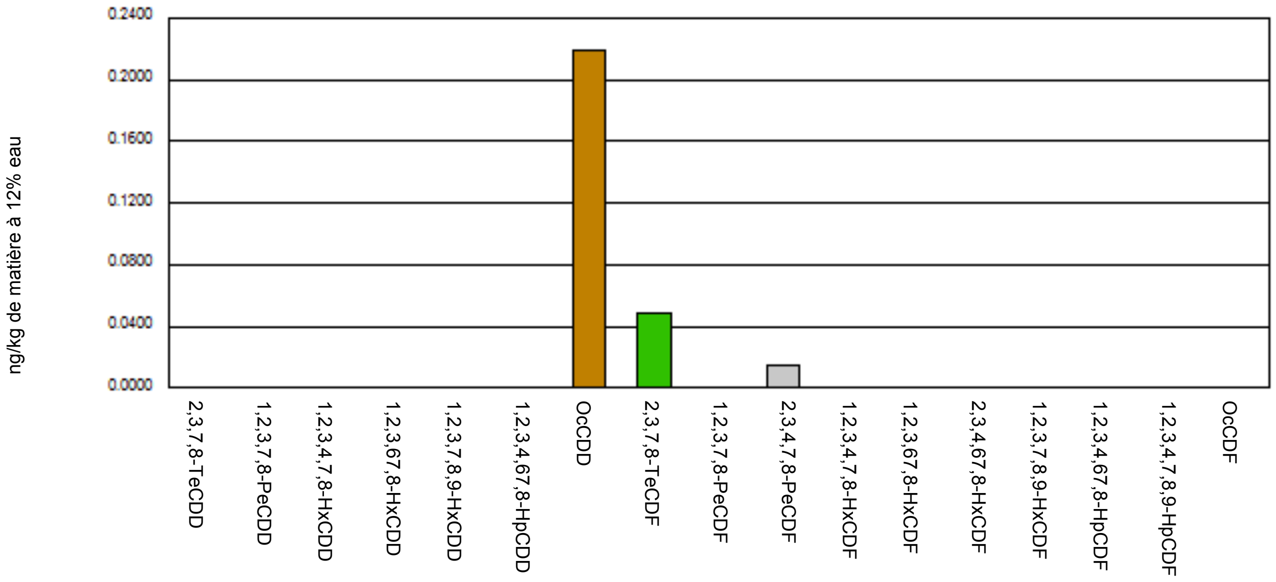


PCDD/F-TEQ des 17 congénères toxiques

Référence CARSO-LSEHL : LSE1708-48971

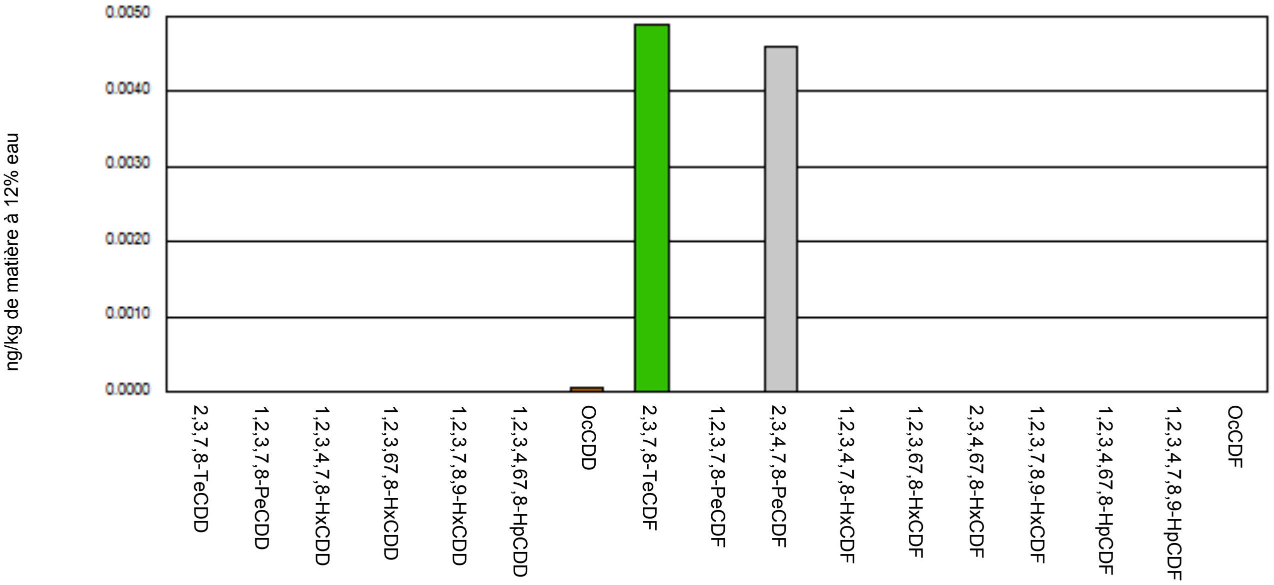




Société par action simplifiée au capital de 2 283 622,30 € - RCS Lyon B 410 545 313 - SIRET 410 545 313 00042 - APE 7120B — N° TVA: FR 82 [illegible]
Siège social et laboratoire : 4, avenue Jean Moulin — CS 30228 - 69633 VENISSIEUX CEDEX - Tél : (33) 04 72 76 16 16 - Fax : (33) 04 78 72 35 03
Site web : www.groupecarso.com -

Institut Scientifique d'Hygiène et d'Analyse
S.A.S au capital de 4 100 000 euros - SIREN : 489 268 045 - APE : 7022 Z
N° TVA intracommunautaire : FR50 489 268 045 - RCS : EVRY 489 268 045
Rue du Chemin Blanc - CHAMPLAN - F-91165 LONGJUMEAU

ISHA
Un laboratoire du Groupe Alpa

**N° Dossier :** MIB.91610
**Code client :** 568020600
**V/ref. :** U8220 V257 LOT: 17207/17209
**Date commande :** 10/08/2017
**Date réception :** 16/08/2017

**SAFE**
**Monsieur BARRAL RENAUD**
**ROUTE DE SAINT BRIS**
**89290 AUGY**

### RAPPORT D'ANALYSE

**Champlan le 24/08/2017**

**Echantillon : K 9187 (MIB.91610.1)**

**Libellé : U8220 V257 CD011344 ENVOI E8795 - REALISER 1 ECH MOYEN**

**Date de prélèvement :**
**Lieu de prélèvement :**
**Température produit :**
**Date préparation :**
**Fournisseur :**
**Date livraison :**
**Date mise en analyse :** 17/08/2017

**Fabriquant :**
**Date Fabr. :**
**DLC :**
**N° de lot :** 17207
**N° agrément :**
**Grammage :** ENV.250G
**Code article :**
**Conditionnement :** 2 POTS
**Mode conservation :**

| Analyse | Méthode | Résultat | Critère | App. | Réf | H/S |
|---|---|---|---|---|---|---|
| * **Microorganismes aérobies 30°C** | NF EN ISO 4833-1 | **< 10 / g** | 100000 | S | Critère demandeur | H |
| **Levures** | Méthode interne (selon NF V08-059) | **< 100 / g** | 1000 | S | Critère demandeur | H |
| **Moisissures** | Méthode interne (selon NF V08-059) | **< 100 / g** | 1000 | S | Critère demandeur | H |
| * **Escherichia coli ß-glucuronidase +** | NF ISO 16649-2 | **< 1 / g** | | | | |
| **Staphylocoques à coagulase positive** | NF EN ISO 6888-3 | **ABSENCE / 1 g** | | | | |
| * **Anaérobies sulfito-réducteurs 46°C (boîtes)** | NF V 08-061 | **< 10 / g** | 100 | S | Critère demandeur | H |
| * **Clostridium perfringens** | NF EN ISO 7937 | **< 1 / g** | | | | |
| **Clostridium perfringens (spores)** | ISHA / CPPRC | **< 1 / g** | | | | |
| * **Bacillus cereus présomptifs** | NF EN ISO 7932 | **< 10 / g** | | | | |
| **Bacillus cereus présomptifs (spores)** | ISHA / BCPRC | **< 10 / g** | | | | |
| * **Salmonella (recherche)** | RBP 31/01-06/08 | **ABSENCE / 25 g** | ABSENCE | S | Critère demandeur | H |
| * **Listeria monocytogenes (recherche)** | AFNOR AES-10/3-09/00 | **ABSENCE / 25 g** | | | | |
| * **Entérobactéries 30°C présumées** | NF V 08-054 | **< 10 / g** | | | | |
| **Pseudomonas** | ISHA / PSENA | **< 10 / g** | | | | |

Stéphanie MAHLER
Responsable d'unité microbiologie

ACCREDITATION
N° 1 -1759
PORTEE
DISPONIBLE SUR
www.cofrac.fr
cofrac
ESSAIS

Ce rapport d'analyse ne concerne que les objets soumis aux analyses. Sa reproduction n'est autorisée que sous forme de fac-similé photographique intégral.
Seuls certains essais rapportés dans ce document sont couverts par l'accréditation de la Section Laboratoire du Cofrac. Ils sont identifiés par le symbole *. Ce rapport comporte 4 pages.

Institut Scientifique d'Hygiène et d'Analyse
S.A.S au capital de 4 100 000 euros - SIREN : 489 268 045 - APE : 7022 Z
N° TVA intracommunautaire : FR50 489 268 045 - RCS : EVRY 489 268 045
Rue du Chemin Blanc - CHAMPLAN - F-91165 LONGJUMEAU



Un laboratoire du Groupe Alpa

**N° Dossier :** MIB.91610
**Code client :** 568020600
**V/ref. :** U8220 V257 LOT: 17207/17209
**Date commande :** 10/08/2017
**Date réception :** 16/08/2017

**SAFE**
**Monsieur BARRAL RENAUD**
**ROUTE DE SAINT BRIS**
**89290 AUGY**

### RAPPORT D'ANALYSE

**Champlan le 24/08/2017**

Appréciation: S=Satisfaisant - A= Acceptable - I=Insatisfaisant / H/S: S=critère de sécurité - H=critère d'hygiène des procédés

Le dossier comporte 2 échantillons

* Analyse accréditée COFRAC. Lorsque tous les paramètres analytiques concernés par la déclaration de conformité sont réalisés sous accréditation, la conclusion globale de conformité est par défaut sous accréditation. La déclaration de conformité ne tient pas compte de l'incertitude de mesure.

**Conclusion critères de sécurité** : *Absence de critères*
**Conclusion hygiène des procédés** : SATISFAISANT

Stéphanie MAHLER
Responsable d'unité microbiologie

ACCREDITATION
N° 1 -1759
PORTEE
DISPONIBLE SUR
www.cofrac.fr

Ce rapport d'analyse ne concerne que les objets soumis aux analyses. Sa reproduction n'est autorisée que sous forme de fac-similé photographique intégral.
Seuls certains essais rapportés dans ce document sont couverts par l'accréditation de la Section Laboratoire du Cofrac. Ils sont identifiés par le symbole *. Ce rapport comporte 4 pages.

Institut Scientifique d'Hygiène et d'Analyse
S.A.S au capital de 4 100 000 euros - SIREN : 489 268 045 - APE : 7022 Z
N° TVA intracommunautaire : FR50 489 268 045 - RCS : EVRY 489 268 045
Rue du Chemin Blanc - CHAMPLAN - F-91165 LONGJUMEAU

ISHA
Un laboratoire du Groupe Alpa

**N° Dossier :** MIB.91610
**Code client :** 568020600
**V/ref. :** U8220 V257 LOT: 17207/17209
**Date commande :** 10/08/2017
**Date réception :** 16/08/2017

**SAFE**
**Monsieur BARRAL RENAUD**
**ROUTE DE SAINT BRIS**
**89290 AUGY**

### RAPPORT D'ANALYSE

**Champlan le 24/08/2017**

**Echantillon : K 9188 (MIB.91610.2)**

**Libellé : U8220 V257 MP ORGANIQUE CD011344 ENVOI E8796 - REALISER 1 ECH MOYEN**

**Date de prélèvement :**
**Lieu de prélèvement :**
**Température produit :**
**Date préparation :**
**Fournisseur :**
**Date livraison :**
**Date mise en analyse :** 17/08/2017

**Fabriquant :**
**Date Fabr. :**
**DLC :**
**N° de lot :** 17209
**N° agrément :**
**Grammage :** ENV.250G
**Code article :**
**Conditionnement :** 2 POTS
**Mode conservation :**

| Analyse | Méthode | Résultat | Critère | App. | Réf | H/S |
|---|---|---|---|---|---|---|
| * **Microorganismes aérobies 30°C** | NF EN ISO 4833-1 | **< 10 / g** | 100000 | S | Critère demandeur | H |
| **Levures** | Méthode interne (selon NF V08-059) | **< 100 / g** | 1000 | S | Critère demandeur | H |
| **Moisissures** | Méthode interne (selon NF V08-059) | **< 100 / g** | 1000 | S | Critère demandeur | H |
| * **Escherichia coli ß-glucuronidase +** | NF ISO 16649-2 | **< 1 / g** | | | | |
| **Staphylocoques à coagulase positive** | NF EN ISO 6888-3 | **ABSENCE / 1 g** | | | | |
| * **Anaérobies sulfito-réducteurs 46°C (boîtes)** | NF V 08-061 | **< 10 / g** | 100 | S | Critère demandeur | H |
| * **Clostridium perfringens** | NF EN ISO 7937 | **< 1 / g** | | | | |
| **Clostridium perfringens (spores)** | ISHA / CPPRC | **< 1 / g** | | | | |
| * **Bacillus cereus présomptifs** | NF EN ISO 7932 | **< 10 / g** | | | | |
| **Bacillus cereus présomptifs (spores)** | ISHA / BCPRC | **< 10 / g** | | | | |
| * **Salmonella (recherche)** | RBP 31/01-06/08 | **ABSENCE / 25 g** | ABSENCE | S | Critère demandeur | H |
| * **Listeria monocytogenes (recherche)** | AFNOR AES-10/3-09/00 | **ABSENCE / 25 g** | | | | |
| * **Entérobactéries 30°C présumées** | NF V 08-054 | **< 10 / g** | | | | |
| **Pseudomonas** | ISHA / PSENA | **< 10 / g** | | | | |

Stéphanie MAHLER
Responsable d'unité microbiologie

ACCREDITATION
N° 1 -1759
PORTEE
DISPONIBLE SUR
www.cofrac.fr
ESSAIS

Ce rapport d'analyse ne concerne que les objets soumis aux analyses. Sa reproduction n'est autorisée que sous forme de fac-similé photographique intégral.
Seuls certains essais rapportés dans ce document sont couverts par l'accréditation de la Section Laboratoire du Cofrac. Ils sont identifiés par le symbole *. Ce rapport comporte 4 pages.

Essai(s) réalisé(s) à l'ISHA : 25, avenue de la république 91300 Massy

**Institut Scientifique d'Hygiène et d'Analyse**
S.A.S au capital de 4 100 000 euros - SIREN : 489 268 045 - APE : 7022 Z
N° TVA intracommunautaire : FR50 489 268 045 - RCS : EVRY 489 268 045
Rue du Chemin Blanc - CHAMPLAN - F-91165 LONGJUMEAU

ISHA
Un laboratoire du Groupe Alpa

**N° Dossier :** MIB.91610
**Code client :** 568020600
**V/ref. :** U8220 V257 LOT: 17207/17209
**Date commande :** 10/08/2017
**Date réception :** 16/08/2017

**SAFE**
**Monsieur BARRAL RENAUD**
**ROUTE DE SAINT BRIS**
**89290 AUGY**

### RAPPORT D'ANALYSE

**Champlan le 24/08/2017**

Appréciation: S=Satisfaisant - A= Acceptable - I=Insatisfaisant / H/S: S=critère de sécurité - H=critère d'hygiène des procédés

Le dossier comporte 2 échantillons

* Analyse accréditée COFRAC. Lorsque tous les paramètres analytiques concernés par la déclaration de conformité sont réalisés sous accréditation, la conclusion globale de conformité est par défaut sous accréditation. La déclaration de conformité ne tient pas compte de l'incertitude de mesure.

**Conclusion critères de sécurité** : *Absence de critères*

**Conclusion hygiène des procédés** : SATISFAISANT

Stéphanie MAHLER

Responsable d'unité microbiologie

ACCREDITATION
N° 1 -1759
PORTEE
DISPONIBLE SUR
www.cofrac.fr

Ce rapport d'analyse ne concerne que les objets soumis aux analyses. Sa reproduction n'est autorisée que sous forme de fac-similé photographique intégral.
Seuls certains essais rapportés dans ce document sont couverts par l'accréditation de la Section Laboratoire du Cofrac. Ils sont identifiés par le symbole *. Ce rapport comporte 4 pages.

SIGMA-ALDRICH

3050 Spruce Street, Saint Louis, MO 63103 USA

### Certificate of Analysis

| | |
|---|---|
| **Product Name:** | AZOXYSTROBIN<br>PESTANAL™, analytical standard |
| **Product Number:** | 31697 |
| **Batch Number:** | BCBT1118V |
| **Brand:** | Sigma-Aldrich |
| **CAS Number:** | 131860-33-8 |
| **Formula:** | $C_{22}H_{17}N_3O_5$ |
| **Formula Weight:** | 403.39 |
| **Expiration Date:** | OCT 2021 |
| **Quality Release Date:** | 04 NOV 2016 |

| TEST | SPECIFICATION | RESULT |
|---|---|---|
| APPEARANCE (COLOR) | WHITE TO YELLOW AND FAINT BEIGE TO LIGHT BEIGE AND FAINT BROWN TO LIGHT BROWN | YELLOW |
| APPEARANCE (FORM) | POWDER OR CRYSTALS | POWDER |
| PURITY (HPLC AREA %) | ≥ 98.0 % | 99.3 % |
| MELTING POINT | 114 - 120 C | 117 C |
| WATER | ≤ 1.0 % | < 0.05% |
| PROTON NMR SPECTRUM | CONFORMS TO STRUCTURE | CONFORMS |

Claudia Geitner

Dr. Claudia Geitner
Manager Quality Control
Buchs, Switzerland

Sigma-Aldrich warrants that at the time of the quality release or subsequent retest date this product conformed to the information contained in this publication. The current specification sheet may be available at Sigma-Aldrich.com. For further inquiries, please contact Technical Service. Purchaser must determine the suitability of the product for its particular use. See reverse side of invoice or packing slip for additional terms and conditions of sale.

#### 31697 Azoxystrobin

| Lot Number: **BCBT1118V** | Sample Name: **T38864_001** |
|---|---|

**Dionex Ultimate 3000 RSLC**

| | | | |
|---|---|---|---|
| *Pump :* | LPG-3400RS | *Injection Time:* | 31.10.16 14:26 |
| *Autosampler:* | WPS-3000 | *Processed By:* | Mikaël Berthet |
| *Detector:* | VWD-3400 | *Vial Number:* | BB4 |
| *Column:* | Ascentis Express C18, 2.7 um | *Column S/N:* | USWM003002 |
| *Column Dim.:* | 150 x 2.1 mm | *Sample Type:* | unknown |
| *Mobile Phase:* | | *Injection Volume:* | 1.0 µl |
| %A : | Acetonitrile | *Flow:* | 0.50 ml/min |
| %B : | Methanol | *Column Temp. (°C):* | 35.0 |
| %C : | Water | *Run Time:* | 15.00 min |
| %D : | Phosphoric acid 0.01M | | |
| *Gradient :* | see Figure 1 | | |
| *Sample Prep.:* | 2 mg sample in 10 ml acetonitrile | | |

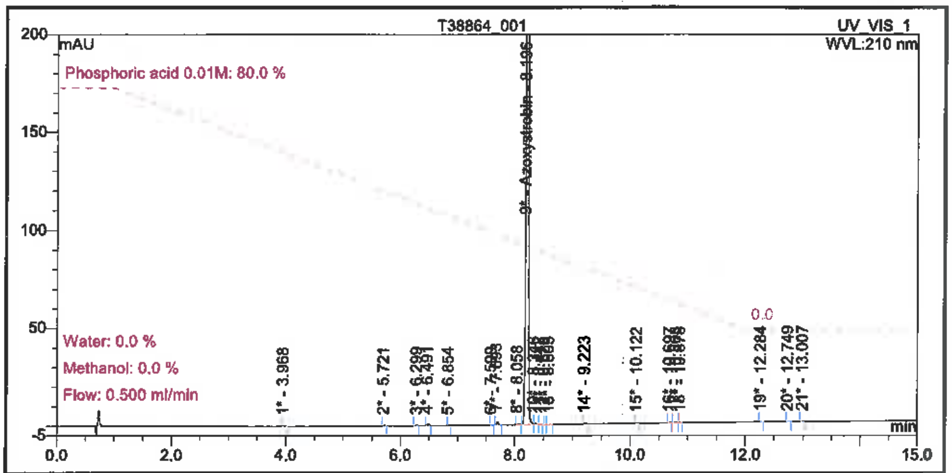


Figure 1: Zoomed Chromatogram

| No. | Ret.Time min | Peak Name | Area mAU*min | Height mAU | Amount | Rel.Area % |
|---|---|---|---|---|---|---|
| 1 | 3.968 | n.a. | 0.01720 | 0.33851 | n.a. | 0.05 |
| 2 | 5.721 | n.a. | 0.00897 | 0.29648 | n.a. | 0.03 |
| 3 | 6.299 | n.a. | 0.00575 | 0.16333 | n.a. | 0.02 |
| 4 | 6.491 | n.a. | 0.02789 | 0.81099 | n.a. | 0.08 |
| 5 | 6.854 | n.a. | 0.00260 | 0.07859 | n.a. | 0.01 |
| 6 | 7.599 | n.a. | 0.00428 | 0.11493 | n.a. | 0.01 |
| 7 | 7.693 | n.a. | 0.05994 | 1.74606 | n.a. | 0.18 |
| 8 | 8.058 | n.a. | 0.01498 | 0.44592 | n.a. | 0.04 |
| 9 | 8.196 | Azoxystrobin | 33.19362 | 967.34672 | n.a. | 99.31 |
| 10 | 8.346 | n.a. | 0.02302 | 0.46552 | n.a. | 0.07 |
| 11 | 8.442 | n.a. | 0.00627 | 0.18528 | n.a. | 0.02 |
| 12 | 8.528 | n.a. | 0.00458 | 0.14295 | n.a. | 0.01 |
| 13 | 8.605 | n.a. | 0.01373 | 0.31659 | n.a. | 0.04 |
| 14 | 9.223 | n.a. | 0.00937 | 0.28877 | n.a. | 0.03 |
| 15 | 10.122 | n.a. | 0.00806 | 0.22493 | n.a. | 0.02 |
| 16 | 10.697 | n.a. | 0.00610 | 0.13251 | n.a. | 0.02 |
| 17 | 10.808 | n.a. | 0.00529 | 0.10469 | n.a. | 0.02 |
| 18 | 10.878 | n.a. | 0.00266 | 0.08675 | n.a. | 0.01 |
| 19 | 12.284 | n.a. | 0.00456 | 0.12120 | n.a. | 0.01 |
| 20 | 12.749 | n.a. | 0.00402 | 0.08758 | n.a. | 0.01 |
| 21 | 13.007 | n.a. | 0.00240 | 0.07375 | n.a. | 0.01 |
| Total: | | | 33.42528 | 973.57205 | | 100.00 |

Table 1: Integration

SIGMA-ALDRICH®

3050 Spruce Street, Saint Louis, MO 63103 USA

### Certificate of Analysis

| | |
|---|---|
| **Product Name:** | BOSCALID<br>PESTANAL™, analytical standard |
| **Product Number:** | 33875 |
| **Batch Number:** | BCBS8868V |
| **Brand:** | Sigma-Aldrich |
| **CAS Number:** | 188425-85-6 |
| **Formula:** | $C_{18}H_{12}Cl_2N_2O$ |
| **Formula Weight:** | 343.21 |
| **Expiration Date:** | AUG 2021 |
| **Quality Release Date:** | 16 SEP 2016 |

| TEST | SPECIFICATION | RESULT |
|---|---|---|
| APPEARANCE (COLOR) | WHITE TO LIGHT YELLOW AND FAINT BEIGE TO LIGHT BEIGE | FAINT YELLOW |
| APPEARANCE (FORM) | POWDER OR CRYSTALS | POWDER |
| PURITY (HPLC AREA %) | ≥ 98.0 % | 99.5 % |
| MELTING POINT | 145 - 150 C | 145 C |
| WATER | ≤ 1.0 % | 0.49 % |
| PROTON NMR SPECTRUM | CONFORMS TO STRUCTURE | CONFORMS |

Claudia Geitner

Dr. Claudia Geitner
Manager Quality Control
Buchs, Switzerland

Sigma-Aldrich warrants that at the time of the quality release or subsequent retest date this product conformed to the information contained in this publication. The current specification sheet may be available at Sigma-Aldrich.com. For further inquiries, please contact Technical Service. Purchaser must determine the suitability of the product for its particular use. See reverse side of invoice or packing slip for additional terms and conditions of sale.

### 33875 Boscalid

| Lot Number: **BCBS8868V** | Sample Name: **00000000560714_001_LC** |
|---|---|

**Dionex Ultimate 3000 RSLC**

| | | | |
|---|---|---|---|
| ***Pump :*** | LPG-3400RS | ***Injection Time:*** | 01.09.16 17:59 |
| ***Autosampler:*** | WPS-3000 | ***Processed By:*** | Mikaël Berthet |
| ***Detector:*** | DAD-3000RS | ***Vial Number:*** | RB6 |
| ***Column:*** | Supelco Ascentis Express C18, 2.7 um | ***Column S/N:*** | - |
| ***Column Dim.:*** | 50 x 2.1 mm | ***Sample Type:*** | unknown |
| ***Mobile Phase:*** | | ***Injection Volume:*** | 0.5 µl |
| %A : | Acetonitrile | ***Flow:*** | 1.00 ml/min |
| %B : | Methanol | ***Column Temp. (°C):*** | 35.0 |
| %C : | Water | ***Run Time:*** | 6.00 min |
| %D : | H3PO4 0.01 M | | |
| ***Gradient :*** | see Figure 1 | | |
| ***Sample Prep.:*** | 0.6 mg sample in 1ml **acetonitrile:water 1:1** | | |

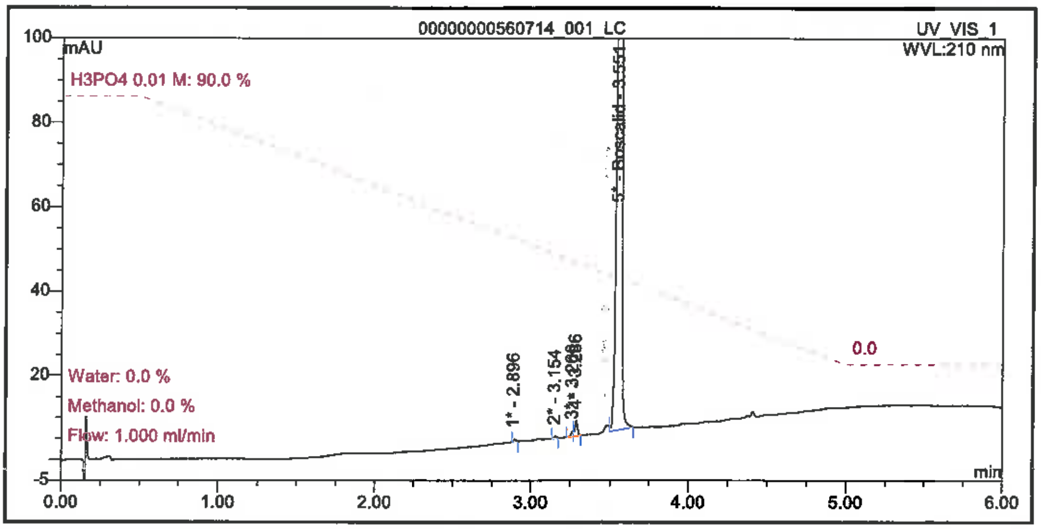


Figure 1: Zoomed Chromatogram

| No. | Ret.Time min | Peak Name | Area mAU*min | Height mAU | Amount | Rel.Area % |
|---|---|---|---|---|---|---|
| 1 | 2.896 | n.a. | 0.01173 | 0.59038 | n.a. | 0.05 |
| 2 | 3.154 | n.a. | 0.00898 | 0.50012 | n.a. | 0.04 |
| 3 | 3.266 | n.a. | 0.02053 | 1.22821 | n.a. | 0.09 |
| 4 | 3.286 | n.a. | 0.06899 | 3.56225 | n.a. | 0.31 |
| 5 | 3.551 | Boscalid | 21.99856 | 1091.96482 | n.a. | 99.50 |
| Total: | | | 22.10881 | 1097.84578 | | 100.00 |

Table 1: Integration

SIGMA-ALDRICH®

3050 Spruce Street, Saint Louis, MO 63103 USA

### Certificate of Analysis

| | |
|---|---|
| **Product Name:** | CHLORPYRIFOS<br>PESTANAL™, analytical standard |
| **Product Number:** | 45395 |
| **Batch Number:** | BCBR6591V |
| **Brand:** | Sigma-Aldrich |
| **CAS Number:** | 2921-88-2 |
| **Formula:** | $C_9H_{11}Cl_3NO_3PS$ |
| **Formula Weight:** | 350.59 |
| **Storage Temperature:** | 2-8 C |
| **Expiration Date:** | JAN 2021 |
| **Quality Release Date:** | 02 MAR 2016 |

| TEST | SPECIFICATION | RESULT |
|---|---|---|
| APPEARANCE (COLOR) | WHITE TO LIGHT YELLOW AND FAINT BEIGE TO LIGHT BEIGE | LIGHT YELLOW |
| APPEARANCE (FORM) | POWDER OR CRYSTALS OR FLAKES OR CHUNKS | SOLID |
| PURITY (HPLC AREA %) | ≥ 98.0 % | 99.3% |
| MELTING POINT | 40 - 46 C | 42C |
| WATER | ≤ 1.0 % | 0.42% |
| PROTON NMR SPECTRUM | CONFORMS TO STRUCTURE | CONFORMS |

Dr. Claudia Geitner
Manager Quality Control
Buchs, Switzerland

Sigma-Aldrich warrants that at the time of the quality release or subsequent retest date this product conformed to the information contained in this publication. The current specification sheet may be available at Sigma-Aldrich.com. For further inquiries, please contact Technical Service. Purchaser must determine the suitability of the product for its particular use. See reverse side of invoice or packing slip for additional terms and conditions of sale.

#### 45395 Chlorpyriphos

**Lot Number: BCBR6591V** | **Sample Name: 00000000541513_001_LC**

**Dionex Ultimate 3000 RSLC**

| | | | |
|---|---|---|---|
| **Pump:** | LPG-3400RS | **Injection Time:** | 26.01.16 16:36 |
| **Autosampler:** | WPS-3000 | **Processed By:** | Mikaël Berthet |
| **Detector:** | DAD-3000RS | **Vial Number:** | GA3 |
| **Column:** | Supelco Ascentis Express C18, 2.7 um | **Column S/N:** | USMD003672 |
| **Column Dim.:** | 50 x 2.1 mm | **Sample Type:** | unknown |
| **Mobile Phase:** | | **Injection Volume:** | 2.0 µl |
| %A : | Acetonitrile | **Flow:** | 1.00 ml/min |
| %B : | Methanol | **Column Temp. (°C):** | 36.0 |
| %C : | Water | **Run Time:** | 6.00 min |
| %D : | Phosphoric acid 0.01 M | | |
| **Gradient:** | see Figure 1 | | |
| **Sample Prep.:** | 1 mg sample in 1 ml acetonitrile:water 1:1 | | |

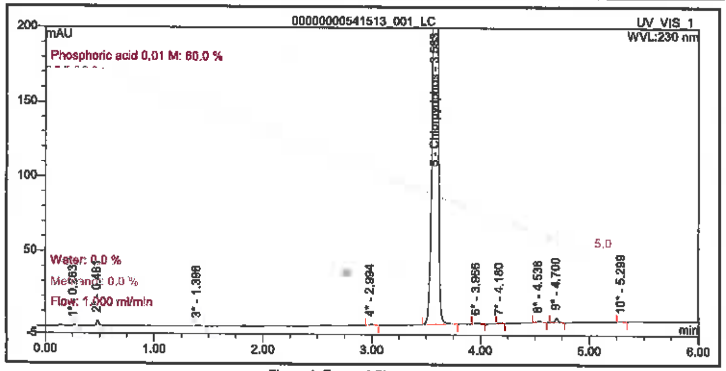


Figure 1: Zoomed Chromatogram

| No. | Ret.Time min | Peak Name | Area mAU*min | Height mAU | Amount | Rel.Area % |
|---|---|---|---|---|---|---|
| 1 | 0.263 | n.a. | 0.00400 | 0.25277 | n.a. | 0.01 |
| 2 | 0.481 | n.a. | 0.07993 | 3.59690 | n.a. | 0.19 |
| 3 | 1.398 | n.a. | 0.01017 | 0.25222 | n.a. | 0.02 |
| 4 | 2.994 | n.a. | 0.02940 | 0.82896 | n.a. | 0.07 |
| 5 | 3.583 | Chlorpyriphos | 42.78613 | 1276.42848 | n.a. | 99.31 |
| 6 | 3.966 | n.a. | 0.02356 | 0.61797 | n.a. | 0.05 |
| 7 | 4.180 | n.a. | 0.01137 | 0.35900 | n.a. | 0.03 |
| 8 | 4.538 | n.a. | 0.03659 | 1.27015 | n.a. | 0.08 |
| 9 | 4.700 | n.a. | 0.09201 | 3.04022 | n.a. | 0.21 |
| 10 | 5.299 | n.a. | 0.00815 | 0.27754 | n.a. | 0.02 |
| Total: | | | 43.08130 | 1286.92421 | | 100.00 |

Table 1: Integration



YB1505.139.fid
Proton_Fluka CDCl3 C:\\ br 39

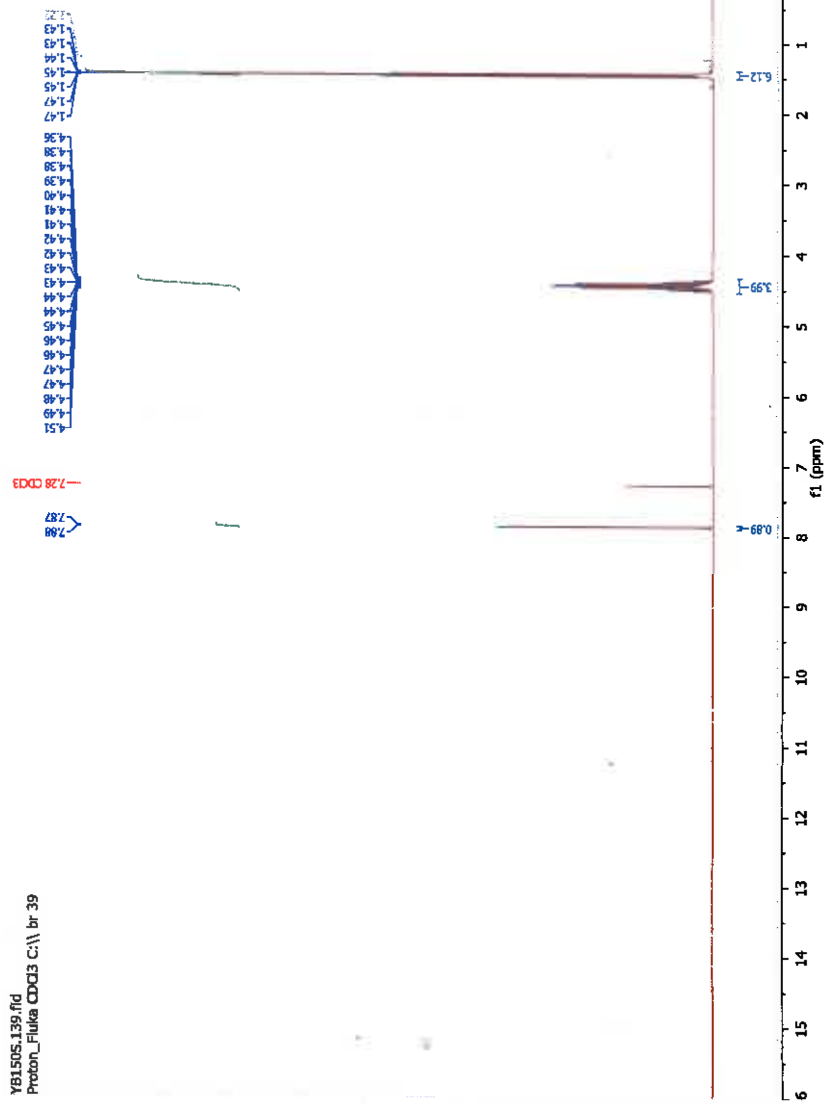

SIGMA-ALDRICH

sigma-aldrich.com

3050 Spruce Street, Saint Louis, MO 63103, USA
Website: www.sigmaaldrich.com
Outside USA:

### Certificate of Analysis

Product Name:
N-(Phosphonomethyl)glycine - 96%

| | |
|---|---|
| **Product Number:** | **337757** |
| **Batch Number:** | **MKCG2949** |
| Brand: | ALDRICH |
| CAS Number: | 1071-83-6 |
| MDL Number: | MFCD00055350 |
| Formula: | C3H8NO5P |
| Formula Weight: | 169.07 g/mol |
| Quality Release Date: | 24 APR 2018 |

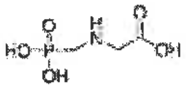

| Test | Specification | Result |
|---|---|---|
| Appearance (Color) | White | White |
| Appearance (Form) | Powder | Powder |
| Infrared Spectrum | Conforms to Structure | Conforms |
| Carbon | 20.3 - 22.3 % | 21.3 % |
| Nitrogen | 7.9 - 8.7 % | 8.2 % |
| Purity (TLC) | $\geq$ 96 % | > 99 % |
| Solubility (Turbidity) 5% in 1N NAOH | Clear to Slightly Hazy | Clear |
| Solubility (Color) | Colorless to Faint Yellow | Colorless |

Michael Grady, Manager
Quality Control
Milwaukee, WI US

Sigma-Aldrich warrants, that at the time of the quality release or subsequent retest date this product conformed to the information contained in this publication. The current Specification sheet may be available at Sigma-Aldrich.com. For further inquiries, please contact Technical Service. Purchaser must determine the suitability of the product for its particular use. See reverse side of invoice or packing slip for additional terms and conditions of sale.

Version Number: 1

SIGMA-ALDRICH®

3050 Spruce Street, Saint Louis, MO 63103 USA

### Certificate of Analysis

| | |
|---|---|
| **Product Name:** | IMIDACLOPRID<br>PESTANAL™, analytical standard |
| **Product Number:** | 37894 |
| **Batch Number:** | BCBT2267 |
| **Brand:** | Sigma-Aldrich |
| **CAS Number:** | 138261-41-3 |
| **Formula:** | $C_9H_{10}ClN_5O_2$ |
| **Formula Weight:** | 255.66 |
| **Expiration Date:** | OCT 2021 |
| **Quality Release Date:** | 22 NOV 2016 |

| TEST | SPECIFICATION | RESULT |
|---|---|---|
| APPEARANCE (COLOR) | WHITE TO LIGHT YELLOW AND FAINT BEIGE TO LIGHT BEIGE | WHITE |
| APPEARANCE (FORM) | POWDER OR CRYSTALS | POWDER |
| PURITY (HPLC AREA %) | ≥ 98.0 % | 100.0 % |
| MELTING POINT | 141 - 146 C | 145 C |
| WATER | ≤ 1.0 % | 0.54 % |
| PROTON NMR SPECTRUM | CONFORMS TO STRUCTURE | CONFORMS |

Dr. Claudia Geitner
Manager Quality Control
Buchs, Switzerland

Sigma-Aldrich warrants that at the time of the quality release or subsequent retest date this product conformed to the information contained in this publication. The current specification sheet may be available at Sigma-Aldrich.com. For further inquiries, please contact Technical Service. Purchaser must determine the suitability of the product for its particular use. See reverse side of invoice or packing slip for additional terms and conditions of sale.

#### 37894 Imidacloprid

Lot Number: **BCBT2267** Sample Name: **T38933_001_LC**

**Dionex Ultimate 3000 RSLC**

***Pump :*** LPG-3400RS
***Autosampler:*** WPS-3000
***Detector:*** VWD-3400
***Column:*** Supelco Ascentis Express C18, 2.7 um
***Column Dim.:*** 50 x 2.1 mm
***Mobile Phase:***
%A : Acetonitrile
%B : Methanol
%C : Water
%D : Phosphoric acid 0.01M
***Gradient :*** see Figure 1
***Sample Prep.:*** 0.5 mg sample in 1ml methanol

***Injection Time:*** 18.11.16 11:17
***Processed By:*** Markus Urthaler
***Vial Number:*** BA2
***Column S/N:*** -
***Sample Type:*** unknown
***Injection Volume:*** 1.0 µl
***Flow:*** 1.00 ml/min
***Column Temp. (°C):*** 35.0
***Run Time:*** 6.00 min

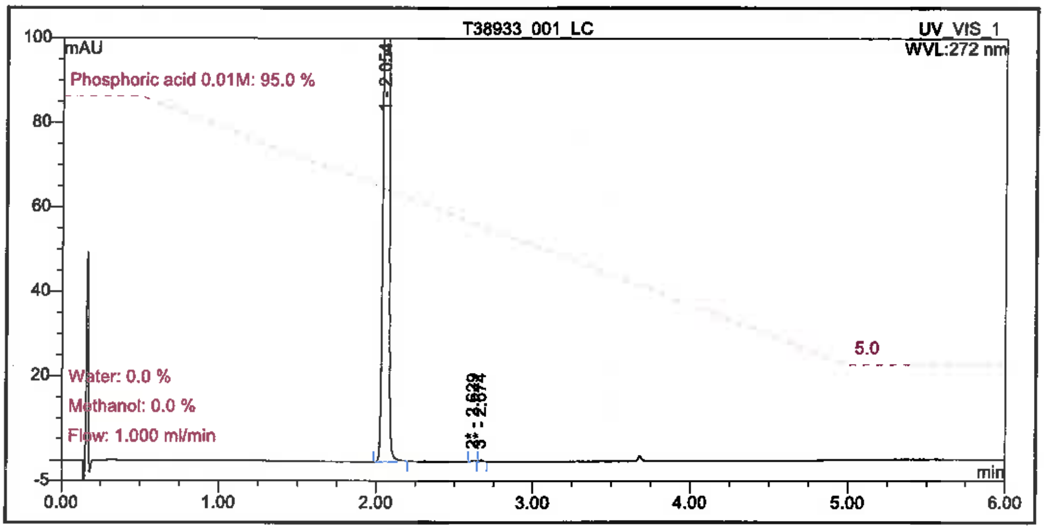


Figure 1: Zoomed Chromatogram

| No. | Ret.Time min | Peak Name | Area mAU*min | Height mAU | Amount | Rel.Area % |
|---|---|---|---|---|---|---|
| 1 | 2.054 | n.a. | 30.01747 | 1460.18351 | n.a. | 99.95 |
| 2 | 2.629 | n.a. | 0.00738 | 0.31070 | n.a. | 0.02 |
| 3 | 2.674 | n.a. | 0.00750 | 0.40892 | n.a. | 0.02 |
| Total: | | | 30.03234 | 1460.90313 | | 100.00 |

Table 1: Integration

SIGMA-ALDRICH®

3050 Spruce Street, Saint Louis, MO 63103 USA

### Certificate of Analysis

| | |
|---|---|
| **Product Name:** | THIABENDAZOLE<br>PESTANAL™, analytical standard |
| **Product Number:** | 45684 |
| **Batch Number:** | BCBV5436 |
| **Brand:** | Sigma-Aldrich |
| **CAS Number:** | 148-79-8 |
| **Formula:** | $C_{10}H_7N_3S$ |
| **Formula Weight:** | 201.25 |
| **Expiration Date:** | JUL 2022 |
| **Quality Release Date:** | 08 AUG 2017 |

| TEST | SPECIFICATION | RESULT |
|---|---|---|
| APPEARANCE (COLOR) | WHITE TO OFF WHITE | WHITE |
| APPEARANCE (FORM) | POWDER OR CRYSTALS | POWDER |
| PURITY (GC AREA %) | ≥ 98.0 % | 98.6 % |
| MELTING POINT | 298 - 304 C | 302 C |
| WATER | ≤ 1.0 % | 0.09 % |
| PROTON NMR SPECTRUM | CONFORMS TO STRUCTURE | CONFORMS |

Claudia Geitner

Dr. Claudia Geitner
Manager Quality Control
Buchs, Switzerland

Sigma-Aldrich warrants that at the time of the quality release or subsequent retest date this product conformed to the information contained in this publication. The current specification sheet may be available at Sigma-Aldrich.com. For further inquiries, please contact Technical Service. Purchaser must determine the suitability of the product for its particular use. See reverse side of invoice or packing slip for additional terms and conditions of sale.

### 45684 Thiabendazole

| Lot Number: **BCBV5436** | Sample Name: **T39427_001_GC** |
|---|---|

**Shimadzu GC-2010 Plus**

| | | | |
|---|---|---|---|
| ***Injector :*** | 250 °C | ***GC Serial Number:*** | C11805270206 US |
| ***Detector (FID):*** | 320 °C | ***Injection Time:*** | 25.07.17 17:02 |
| ***Carriergas:*** | Helium | ***Sample Type:*** | unknown |
| ***Velocity:*** | 40.0 cm/s | ***Channel:*** | GC_2 |
| ***Temp. Control:*** | see Figure 1 | ***Sequence Creation Time:*** | 04.07.17 15:51 |
| ***Split Ratio:*** | 100:1 | ***Sequence Created By:*** | Mirela Brdanovic |
| ***Range:*** | - | ***Time Processed:*** | 11.08.17 09:14 |
| ***Injection Volume:*** | 1.0 ul | ***Processed By:*** | Markus Urthaler |
| ***Vial Number:*** | 4 | | |
| ***Run Time:*** | 20.00 min | | |
| ***Column:*** | Equity-1, 30 m x 0.25 mm, 1.0 um | | |
| ***Sample Prep.:*** | ca 20mg sample in 2ml methanol | | |

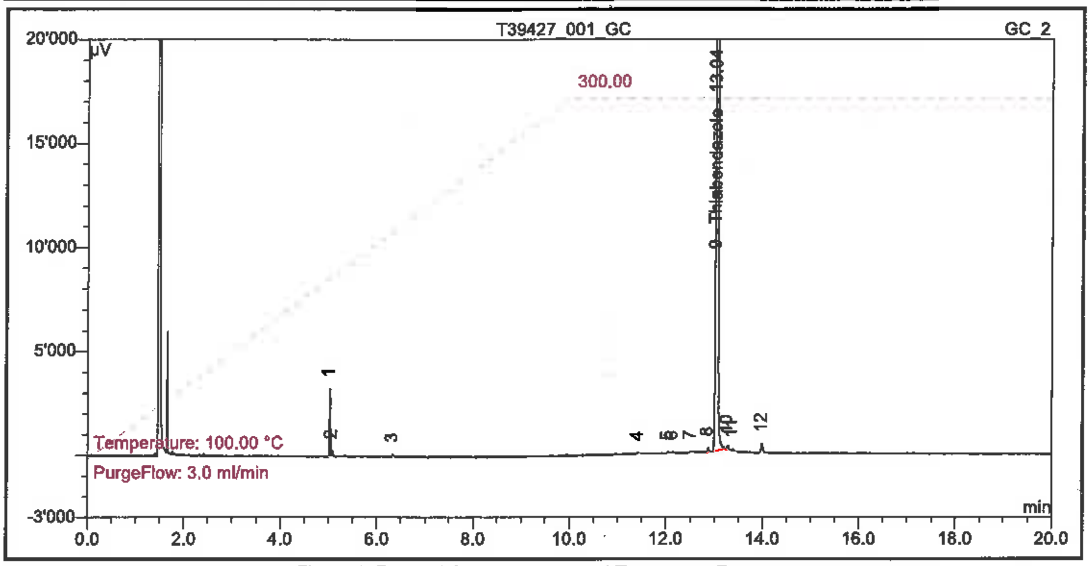


Figure 1: Zoomed Chromatogram and Temperatur Program

| No. | Ret.Time min | Peak Name | Type | Height µV | Area µV*min | Amount | Rel.Area % |
|---|---|---|---|---|---|---|---|
| 1 | 5.028 | n.a. | BM | 3257 | 59.4046 | n.a. | 0.714 |
| 2 | 5.074 | n.a. | MB | 269 | 5.5508 | n.a. | 0.067 |
| 3 | 6.325 | n.a. | BMB | 115 | 1.7783 | n.a. | 0.021 |
| 4 | 11.430 | n.a. | BMB* | 80 | 2.8125 | n.a. | 0.034 |
| 5 | 12.042 | n.a. | BMB* | 76 | 1.9471 | n.a. | 0.023 |
| 6 | 12.143 | n.a. | BMB* | 88 | 3.4967 | n.a. | 0.042 |
| 7 | 12.516 | n.a. | BMB* | 83 | 3.0967 | n.a. | 0.037 |
| 8 | 12.871 | n.a. | BMB* | 210 | 7.2075 | n.a. | 0.087 |
| 9 | 13.040 | Thiabendazole | BMB | 245305 | 8210.3329 | n.a. | 98.619 |
| 10 | 13.274 | n.a. | BMB | 204 | 6.9992 | n.a. | 0.084 |
| 11 | 13.377 | n.a. | BMB* | 108 | 6.3367 | n.a. | 0.076 |
| 12 | 13.988 | n.a. | BMB | 423 | 16.3450 | n.a. | 0.196 |
| Total: | | | | 250218 | 8325.3079 | | 100.000 |
