## Supplementary Material 2. Shotgun metagenomics analysis at the phylum level. for "Shotgun metagenomics and metabolomics reveal glyphosate alters the gut microbiome of Sprague-Dawley rats by inhibiting the shikimate pathway"

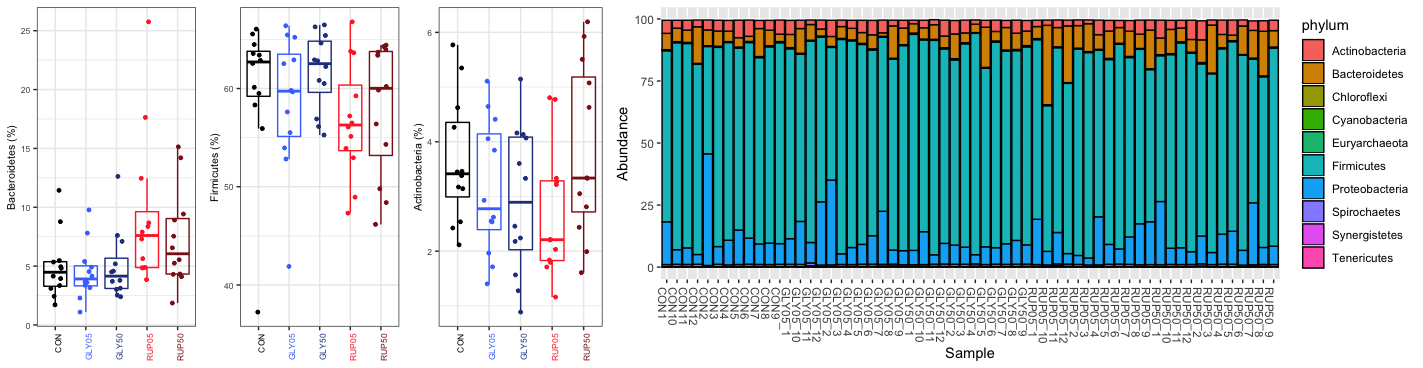

**Supplementary Material 2. Shotgun metagenomics analysis at the phylum level. A.** Scatter plots displaying individual changes in relative abundance for 3 majo phyla. **B**.Top 10 phyla in the rat caecal microbiome based on the results of the comparison to the BGI gut microbiome catalogue
